# Intracellular compartmentalization shapes lipid access and metabolic fitness of mycobacteria

**DOI:** 10.64898/2026.02.10.704997

**Authors:** Mélanie Foulon, Sylvana V. Hüttel, Leonhard Breitsprecher, Wiebke Gauda, Lyudmil Raykov, Hendrik Koliwer-Brandl, Miriam Ohlhagen, Roland Thünauer, Dominik Schwudke, Hubert Hilbi, Thierry Soldati, Caroline Barisch

## Abstract

Intracellular mycobacteria encounter distinct metabolic environments as they transition between vacuolar and cytosolic compartments within host cells, yet how nutrient access is shaped by this compartmentalization remains poorly understood. Here, we use *Mycobacterium marinum* fatty acyl– CoA ligase 6 (FACL6) as a functional entry point to examine how lipid acquisition and processing are coordinated during intracellular infection. By combining host and bacterial genetic perturbations with dual RNA-sequencing and high-resolution imaging in genetically tractable amoebal infection models, and validating key phenotypes in mammalian cells, we show that lipid metabolic programs in intracellular mycobacteria are tightly linked to subcellular localization. Sterol utilization and neutral lipid storage are preferentially engaged during the intravacuolar phase, whereas cytosolic exposure is associated with reduced lipid accumulation. Deletion of *facl6* disrupts this coordinated scenario, resulting in altered bacterial cell envelope architecture, premature membrane damage, defective neutral lipid storage, and reduced intracellular fitness despite enhanced cytosolic access. Together, these findings reveal that loss of FACL6 causes intrinsic defects in lipid handling and highlight how compartment-specific lipid environments shape the outcome of mycobacterial infection.

**Teaser:** Loss of FACL6 reveals how intracellular compartmentalization shapes mycobacterial lipid metabolism

## Introduction

Tuberculosis (Tb) remains one of the leading infectious causes of death worldwide, driven by the ability of *Mycobacterium tuberculosis* (Mtb) to persist within host cells and organs. The nutritional status of the host and particularly undernourishment or starvation exacerbates disease progression, while improved nutrition has been associated with better outcomes, highlighting the importance of nutrient availability for both, the host, and the pathogen. These nutritional states might ultimately shape the local microenvironment within granulomas, which serve as both, immunological barriers, and metabolically specialized niches that Mtb must adapt to. Granulomas, the hallmark of Mtb infections, are organised and highly heterogeneous structures. Counteracting its established immune function, this environment is increasingly recognised as nutrient source for mycobacteria for which the accumulation of lipid droplet (LD)-laden macrophages is characteristic. Lipids (i.e. fatty acids and sterols) released from LDs and membranes (Barisch and Soldati, 2017) might support intracellular mycobacterial replication and persistence (Simwela et al., 2025). These lipids are used (i) to generate energy by β-oxidation, (ii) as building blocks for cell wall lipids, and (iii) for the formation of intracellular lipid inclusions (ILIs) (Barisch et al., 2015b, Barisch and Soldati, 2017).

*Mycobacterium marinum* (Mm) is a widely used model to study Mtb infection, as it shares conserved virulence factors, metabolic pathways, and an intracellular lifestyle with Mtb, while offering experimental advantages such as faster growth and genetically tractable host models (for a review, see (Guallar-Garrido and Soldati, 2024)). Importantly, Mm is known to induce Tb-like diseases, with granulomatous lesions, in ectothermic hosts such as zebrafish and frogs. Based on extensive experimental evidence using both Mtb and Mm, a conserved model for an intracellular infection cycle has emerged (Guallar-Garrido and Soldati, 2024, Cardenal-Munoz et al., 2017b). Following phagocytic uptake, mycobacteria initially interfere with the phagocytic pathway and reside within a *Mycobacterium*-containing vacuole (MCV). Subsequent ESX-1-dependent damage to the vacuolar membrane triggers host membrane repair responses, resulting in a dynamic damage-repair cycle. These responses involve ‘endosomal sorting complex required for transport’ (ESCRT)-mediated membrane repair, autophagy-related pathways (Cardenal-Munoz et al., 2017a), which together limit bacterial cytosolic access and intracellular replication, while ER-dependent membrane repair provides lipids for membrane restoration (Anand et al., 2023). In line with this, ESX-1-deficient (RD1) mutants largely remain confined within vacuolar compartments, whereas Mm wild-type (wt) escapes such restriction in autophagy-deficient (*atg1*-KO) host cells (Cardenal-Munoz et al., 2017a, Bernard et al., 2020, López-Jiménez et al., 2018). Moreover, the phosphatases PtpA, PtpB, and SapM promote the escape of Mm from the pathogen vacuole to the cytoplasm (Koliwer-Brandl et al., 2019). Persistent MCV membrane disruption allows bacteria to ultimately access the host cytosol before disseminating to other cells. Throughout this cycle, mycobacteria encounter distinct intracellular compartments with potentially differing nutrient availability, underscoring the need for metabolic plasticity during infection. While the availability of nutrients such as iron limits intracellular growth of Mm (Knobloch et al., 2020), the contribution of nutrient availability in different intracellular compartments has not been assessed.

Given the centrality of host-derived fatty acids for intracellular survival, mycobacteria possess a very diverse set of lipid-modifying enzymes and transport systems to access and import host lipids. Secreted lipases and esterases hydrolyze host triglycerides, cholesteryl esters, and membrane phospholipids, releasing fatty acids that support intracellular growth and virulence (Lin et al., 2024). These lipids are then imported through dedicated multiprotein transporters, including the Mammalian Cell Entry (Mce) complexes Mce1 and Mce4, which mediate uptake of fatty acids and cholesterol, respectively, with additional coordination provided by factors such as LucA (Pandey and Sassetti, 2008, Nazarova et al., 2019, Nazarova et al., 2017). Then, fatty acyl-CoA synthetases (FACLs) convert free fatty acids into acyl-CoA thioesters for catabolism and storage, and fatty acyl-AMP ligases (FAALs) generate acyl-AMP intermediates that are redirected into complex cell-envelope lipid biosynthesis (Trivedi et al., 2004, Mohanty et al., 2011).

FACL6 is an acyl-CoA synthetase that is highly conserved between Mtb and Mm. It belongs to the conserved fatty acid transport protein (‘FATP‘) family found across all kingdoms of life and has been implicated in fatty acid uptake and activation. This process, known as ‘vectorial acylation‘, generates acyl-CoAs that are effectively trapped within the cell or its subcellular compartments, thereby driving continued fatty-acid uptake by maintaining a concentration gradient. Strikingly, Mtb encodes 34 fatty acyl-CoA synthetases (Fad) D-type adenylating enzymes (Duckworth et al., 2012), compared to only one in most bacterial species, highlighting the complexity and specialization of fatty acid activation in this pathogen. In line with that, expression of FACL6 (Rv1206/FadD6 in Mtb) in COS cells and *E. coli* facilitated the uptake of free fatty acids, suggesting a potential role in lipid import (Daniel et al., 2014, Hirsch et al., 1998, Mallick et al., 2021). FACL6 itself is a peripheral membrane protein with broad substrate specificity, capable of activating a range of saturated and unsaturated long-chain fatty acids (Mater et al., 2022). Genetic disruption of *facl6* in Mtb impaired incorporation of radiolabeled fatty acids and reduced triacylglycerol (TG) levels during *in vitro* dormancy, linking FACL6 to lipid metabolism under stress conditions (**Fig. 1A**; (Daniel et al., 2014)). Finally, FACL6 has been proposed as a peripheral membrane-associated enzyme with broad substrate specificity (Fig. 1A; (Daniel et al., 2014, Mater et al., 2022)). However, since fatty acid activation in mycobacteria supports not only energy production and storage but also the biosynthesis of cell envelope lipids, including mycolic acids and other virulence-associated lipids such as phthiocerol dimycocerosate (PDIM) or phenolic glycolipids (PGLs) (Trivedi et al., 2004, Mohanty et al., 2011), FACL6 occupies a central position in lipid metabolic networks. In line with this, *in silico* analysis of ILI-associated proteomes supports the notion that FACL6 is part of a core of enzymes, which synthesize CoA esters of fatty acids released by TG hydrolysis (**Fig. 1A**; (Dargham et al., 2023). Together, these findings place FACL6 at the interface of multiple fatty acid pools within the bacterial cell.

**Figure 1.**
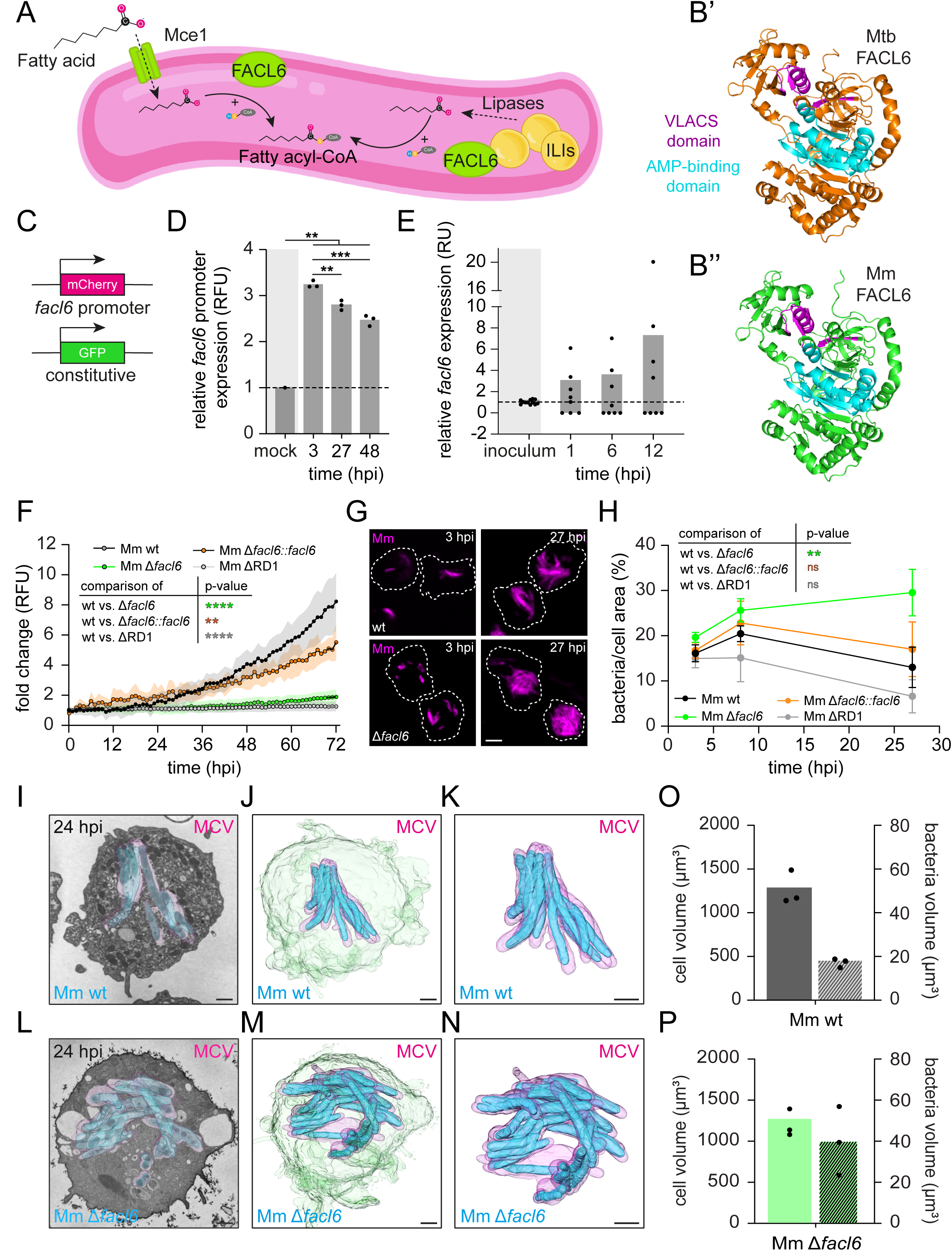
Mm. Δ***facl6* has an intracellular growth defect and is forming microcolonies.** (**A**) Schematic representation of the central role of FACL6 in exogenous and endogenous fatty acid activation in mycobacteria. (**B**) AlphaFold predicted 3D protein structures of Mtb (**B’**) and Mm FACL6 (**B’’**), showing the conserved AMP-binding and the very long-chain acyl-CoA synthetase (VLACS) domains. (**C**) Schematic representation of the dual-fluorescence reporter strain used to monitor *facl6* promoter activity. (**D**) Relative *facl6* promoter expression during Dd infection measured by flow cytometry at 3, 27 and 48 hpi (**E**) *facl6* relative expression (to *gapdh*) during Dd infection measured by RT-qPCR at 1, 6 and 12 hpi. (**F**) Intracellular growth of Mm wt, Δ*facl6*, Δ*facl6::facl6* or ΔRD1 expressing GFP reporter in Dd was followed by microplate reader over 72 hours. Growth curves show mean fold change ± SD (N=3) analyzed using a two-way ANOVA with Fisher’s LSD test (**p<0.01, ****p<0.0001). RFU: Relative Fluorescence Unit. (**G**-**H**) Intracellular bacteria were observed by spinning disc microscopy at 3 and 27 hpi (scale bars, 5 µm), and the area occupied by the bacteria inside each cell was measured and quantified for various Mm strains (mean ± SEM, N=3, statistical differences were determined using an unpaired two-tailed Student’s t-test, **p<0.01; ns: not significant). (**I-N**) 3D reconstruction of Dd cells infected with Mm wt (**I-K**) or Mm Δ*facl6* (**L-N**) at 24 hpi, using serial block-face scanning electron microscopy (SBF-SEM). (**O**-**P**) Quantifications of the volumes occupied by Mm wt (**O**) or Mm Δ*facl6* (**P**) bacteria within the infected cells from SBF-SEM 3D reconstructions. Each dot represents a single cell from two independent biological replicates.

More generally, fatty acid-derived metabolites can be funneled into distinct metabolic routes depending on cellular demands and nutrient availability. For example, propionyl-CoA, resulting from catabolism of sterols and odd-chain fatty acids can be used to fuel the methylcitrate cycle (MCC) or be detoxified via the methylmalonyl-CoA pathway (MMP) pathway for the synthesis of cell wall lipids (VanderVen et al., 2015, Griffin et al., 2012). However, despite the demonstrated role of FACL6 in lipid metabolism under *in vitro* conditions, there is currently no evidence that FACL6 is directly associated with Mce1-mediated fatty acid uptake or with TG turnover in the context of infection.

Although intracellular mycobacteria accumulate lipids in form of ILIs, the extent to which these structures reflect active nutrient acquisition versus storage remains poorly understood. This is further complicated by the compartmental context - i.e. whether the bacteria reside in a phagosome-derived compartment, the MCV, or the cytosol - each presenting a distinct metabolic environment. While intracellular mycobacteria seem to be well equipped to access host-derived lipids, it remains unclear which specific lipid species are actually taken up, when, and under which intracellular conditions. Nutrient availability likely differs depending on the compartment. In the MCV, the environment is rich in membranes and sterols as well as LDs, which somehow translocate into the vacuolar lumen, potentially providing access to TGs (Barisch et al., 2015b, Peyron et al., 2008). In contrast, the cytosol is considered poor in free-lipids, and is enriched in sugars and amino acids; nutrient classes that are non-typical carbon sources for intracellular mycobacteria. Notably, interactions between cytosolic mycobacteria and LDs have been observed (Barisch et al., 2015b), raising the question about the accessibility and relevance of LD-derived lipids for mycobacteria outside their MCV.

To determine how nutrient access differs between intracellular compartments, we combined host and bacterial genetic perturbations across multiple infection models and used FACL6 as a functional entry point to probe lipid acquisition by intracellular mycobacteria. Importantly, Mtb and Mm FACL6 share 83.08 % identity and both contain the conserved AMP-binding domain and the very long-chain acyl-CoA synthetase (VLACS) domains (**Fig 1B**). As host systems, we used *Dictyostelium discoideum* (Dd), BV-2 mouse microglial cells (BV-2) and immortalized bone marrow-derived macrophages (iBMDMs) from C57BL/6 mice (Broz et al., 2010, De Nardo et al., 2018), a commonly used mouse strain with well-characterized immune responses. While Dd is genetically tractable and extensively characterized in terms of lipid metabolism and infection biology (Guallar-Garrido and Soldati, 2024, Dunn et al., 2017), it is also relevant for imaging mycobacterial infections (Barisch et al., 2015a, Franzkoch et al., 2024). BV-2 cells are particularly well suited for high-content imaging (Trofimov et al., 2018) and are relevant for extra-pulmonary TB research, as Tb meningitis represents the deadliest form of the disease (Tram et al., 2025). Finally, immortalized bone marrow– derived macrophages (iBMDMs) provide a robust system for studying infection-induced cell death, especially in the context of host autophagy (Pruenster et al., 2023). By integrating dual RNA-sequencing to distinguish transcriptional programs of cytosolic versus vacuolar bacteria with serial block-face electron microscopy (SBF-SEM) to assess intracellular lipid inclusions in a compartment-specific manner, we examined how lipid availability and bacterial adaptation are shaped by subcellular localization. The combination of models and approaches further enabled us to assess how the absence of FACL6 affects bacterial organization into intracellular microcolonies and is associated with imbalanced lipid metabolism and alterations of the bacterial cell surface. Altogether, this work allowed us to assess how access to host-derived lipids varies across intracellular niches and how mycobacteria adjust their metabolic program accordingly.

## Results

### FACL6 is expressed throughout intracellular infection

To determine when FACL6 is expressed during infection, we first monitored *facl6* transcription using a dual-fluorescence reporter strain of Mm expressing GFP constitutively and mCherry under the control of the *facl6* promoter (**Fig. 1C**). This flow cytometry-based approach allowed quantitative assessment of promoter activity at defined early, intermediate, and late stages of infection (3-, 27-, and 48-hours post-infection (hpi), respectively). The *facl6* promoter activity, reflected by its expression relative to GFP constitutive expression, was detected throughout the infection time course, with maximal activity at 3 hpi followed by a progressive decline at later time points (**Fig. 1D**). To complement this reporter-based analysis and gain higher temporal resolution during the early phase of infection, we next quantified *facl6* expression by RT-qPCR in intracellular bacteria, using broth-grown Mm (i.e. inoculum) as a reference. This analysis revealed an heterogenous expression of *facl6* during infection, with maximal transcript levels appearing highest around 12 hpi (**Fig. 1E**), corresponding to the damage-repair phase of the MCV. Together, these two independent approaches indicate that *facl6* is induced early during intracellular infection and remains expressed, albeit at decreasing levels, during later stages.

### Deletion of *facl6* causes intracellular growth attenuation and microcolony formation

To assess the functional role of FACL6 during infection, we generated a deletion mutant of *facl6* (MMAR_4232) and a complemented strain expressing *facl6* from an integrative plasmid (pMV361) under the control of the *hsp60* promoter. Loss and partial restoration of *facl6* expression were confirmed by qRT-PCR (**Fig. S1A**). Deletion of *facl6* did not affect bacteria uptake by host cells, allowing direct comparison of intracellular growth kinetics (**Fig. S1B**). Across multiple phagocytic hosts, including Dd, BV-2 murine microglial cells, and iBMDMs, the Mm Δ*facl6* mutant displayed a pronounced intracellular growth defect compared to Mm wt bacteria and the complemented strain (**Fig. 1F; Fig. S1C-E**). This attenuation was consistently observed throughout the infection time course. Live-cell imaging revealed that intracellular Mm Δ*facl6* bacteria frequently formed distinct microcolonies in all host cell types examined (**Fig. 1G-H; Fig. S1D and F**). Quantitative image analysis showed that the segmented bacteria area per infected cell was significantly larger for the Mm Δ*facl6* mutant than for Mm wt bacteria, despite reduced overall bacteria proliferation (**Fig. 1H)**. In contrast, the ESX-1-deficient Mm ΔRD1 mutant, which remains confined within an intact MCV, formed smaller bacteria clusters throughout infection, while the complemented Mm Δ*facl6::facl6* strain closely resembled the Mm wt phenotype. To characterize these microcolonies at higher resolution, we performed SBF-SEM. This analysis revealed that Mm Δ*facl6* microcolonies consist of densely packed bacteria with irregular orientation, in contrast to the more ordered bacterial assemblies observed for Mm wt (**Fig. 1I-P**). Volumetric reconstruction suggested that Mm Δ*facl6* microcolonies were larger than those formed by Mm wt bacteria (**Fig. 1O-P**). Together, these results indicate that absence of FACL6 leads to a marked intracellular growth defect accompanied by the formation of enlarged, disorganized microcolonies, indicating a critical role for FACL6 in regulating bacteria proliferation and spatial organization during intracellular infection.

### Cytosolic access is not sufficient to restore intracellular growth of Mm Δ*facl6*

To determine whether the intracellular growth defect and microcolony phenotype of the Mm Δ*facl6* mutant are linked to altered vacuolar confinement or limited access to the host cytosol, we next examined its behavior in host cells with impaired membrane repair and xenophagic restriction, i.e. a host cell defense pathway targeting cytoplasmic pathogens including mycobacteria (Deretic et al., 2013, Aylan et al., 2023, Cardenal-Munoz et al., 2017a). The early stage of Mm infection is shaped by the dynamic interplay between bacterial membrane-damaging activities and host-mediated repair of the MCV, which ultimately determines bacterial confinement or access to the cytosol. Disruption of host sensing and repair pathways, such as in Dd *atg1*-KO cells, results in defective membrane repair, premature escape of bacteria into the cytosol, and impaired restriction of cytosolic bacteria by xenophagy (**Fig. 2A**; (Cardenal-Munoz et al., 2017a)). This genetic background therefore allows discrimination between “passive” attenuation resulting from confinement to the MCV and attenuation caused by “active” restriction within the cytosol. Accordingly, Mm wt induces MCV damage and proliferates more extensively in the cytosol of Dd *atg1*-KO cells than in Dd wt cells (**Fig. 2A)**, whereas the ESX-1 deficient Mm ΔRD1 mutant does not induce vacuolar damage, fails to access the cytosol, and remains confined in either Dd strain (Cardenal-Munoz et al., 2017a).

**Figure 2.**
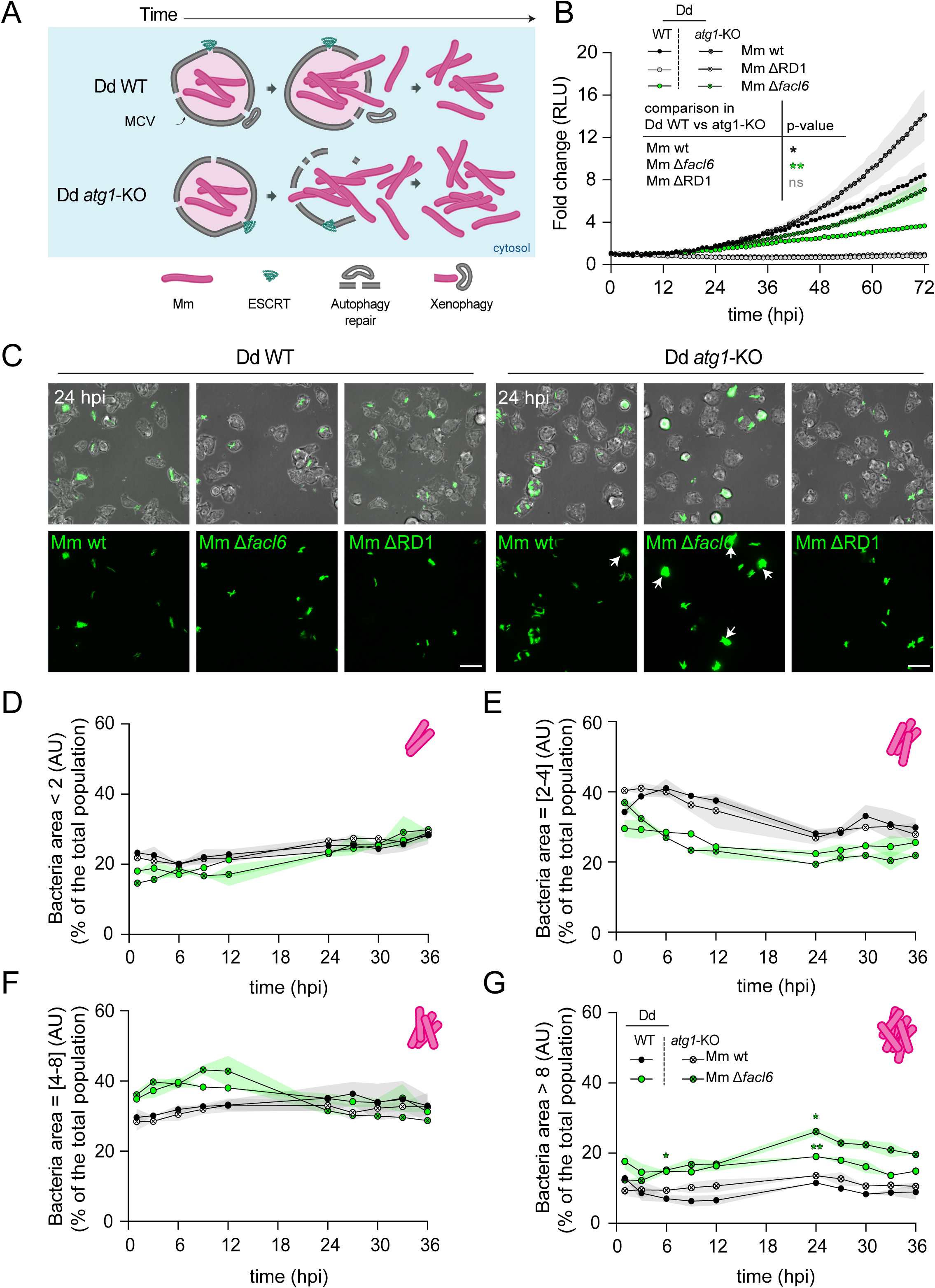
Mm. Δ***facl6* intracellular growth is partially restored in absence of autophagy.** (**A**) Schematic representation of Mm wt fate in Dd wt or *atg1*-KO over the time-course of the infection. **(B)** Intracellular growth of Mm in Dd wt or *atg1*-KO. Cells were infected with luminescent Mm wt, ΔRD1 or Δ*facl6.* Luminescence was measured for 72 hours using a microplate reader. Growth curves show mean fold change ± SEM (N=3) analyzed using a two-way ANOVA with Fisher’s LSD test (*p<0.05, **p<0.01. RLU: Relative Luminescence Unit. (**C)** High-content images illustrating microcolony formation (arrows). Dd WT or *atg1*-KO cells were infected with Mm wt, ΔRD1 or Δ*facl6* expressing GFP and observed by spinning disc microscopy at 24 hpi. (scale bars, 10 µm). (**D-E**) Quantification of the proportions of bacterial events with an area < 2 AU (**D**), in between 2 and 4 AU (**E**), in between 4 and 8 (**F**) and > 8 AU (**G**) (as a percentage of total bacterial events/population) over the time-course of the infection by high-content microscopy for 36 hpi.

Strikingly, the Mm Δ*facl6* mutant exhibited enhanced intracellular growth in Dd *atg1*-KO cells compared to Dd wt cells, reaching bacterial loads comparable to Mm wt in Dd wt (**Fig. 2B**). This increase indicates that loss of *facl6* promotes membrane damage and cytosolic access, conferring a growth advantage when host repair and xenophagic restriction are impaired. Notably, along with enhanced cytosolic access, Mm Δ*facl6* maintained a pronounced microcolony growth pattern in both Dd wt and Dd *atg1*-KO cells (**Fig. 2C**), a phenotype that was also observed in BV-2 macrophages (**Fig. S2A-B**). Quantitative classification of intracellular bacterial events based on segmented area into different size-defined populations further supported these observations, revealing a progressive shift toward larger bacterial areas for Mm Δ*facl6*, both in Dd wt and atg1-KO cells, throughout the infection **(Fig. 2D-G)**. However, even under conditions favoring sustained cytosolic residence in Dd *atg1*-KO cells, Mm Δ*facl6* did not proliferate to the same extent as Mm wt under these conditions, indicating that cytosolic localization alone is not sufficient to allow unrestricted intracellular growth.

### Dual RNA sequencing reveals compartment- and time-dependent host-pathogen transcriptional responses during Mm Δ*facl6* infection

To investigate the molecular basis underlying this phenotype, we next characterized host and bacterial transcriptional responses during infection using dual RNA-sequencing (RNA-seq). We first compared the global transcriptional profiles of Dd wt and *atg1*-KO cells infected with Mm wt or Mm Δ*facl6* across infection conditions. Host transcriptional responses were broadly similar between infections with the two bacterial strains in both genetic backgrounds (**Fig. 3A**). However, a pronounced divergence emerged at 6 hpi, indicating that this time point represents a critical stage at which host responses to Mm wt and Mm Δ*facl6* are most visible. This early divergence suggested that host cells detect qualitative or quantitative differences in the interaction with Mm Δ*facl6* well before differences in bacterial burden become apparent.

**Figure 3.**
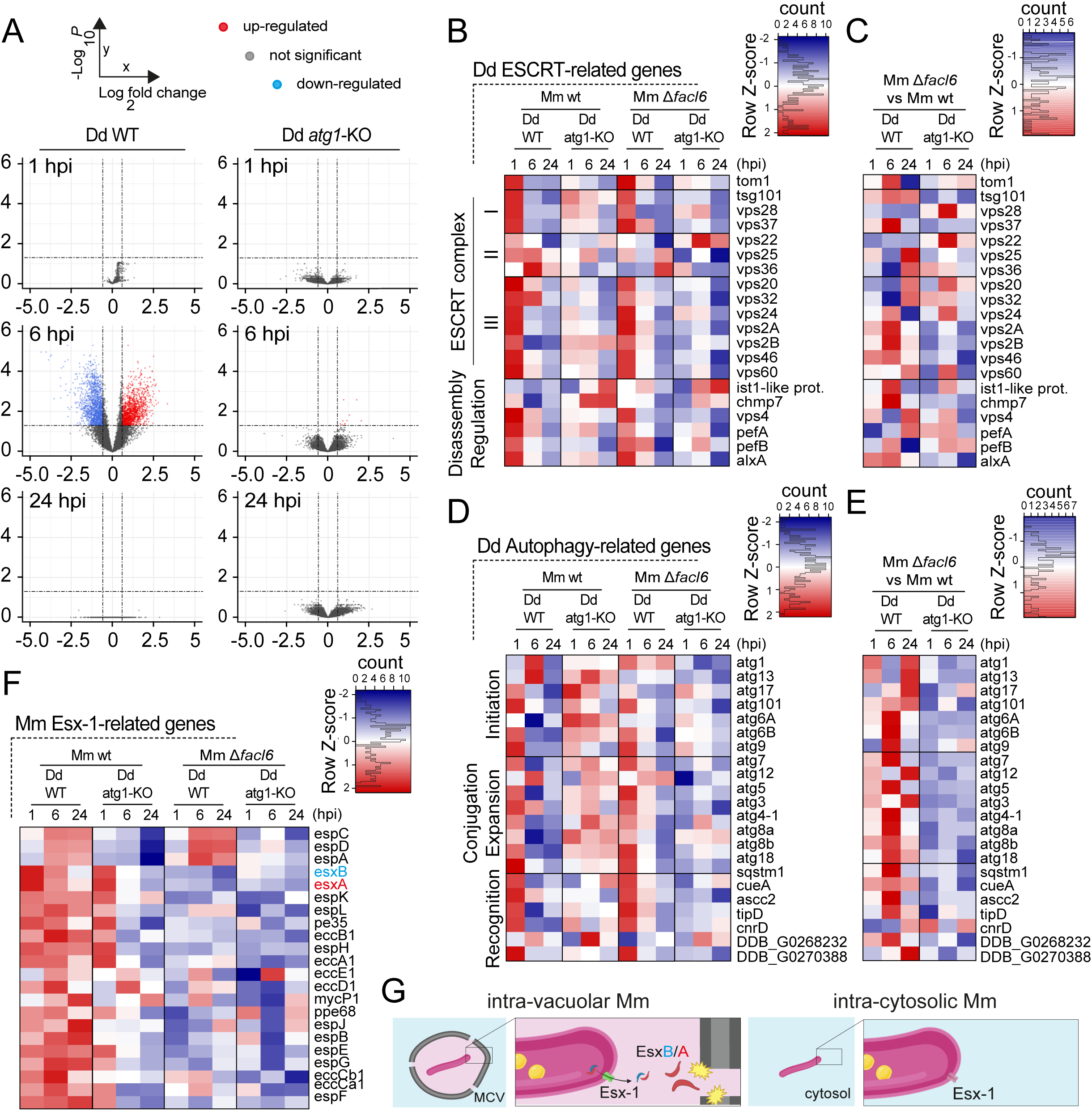
Transcriptomic signature of cells infected with Mm. Δ***facl6* highlights a more sustained host membrane damage-response.** Temporal dual RNAseq analysis of Dd WT or *atg1*-KO FACS-sorted cells infected with GFP-expressing Mm wt or Δ*facl6.* Total RNA was extracted at 1, 6 and 24 hpi. Dual-RNA-sequence libraries were prepared after rRNA depletion and sequenced; reads were aligned separately to the Dd and Mm reference genomes. (**A**) Volcano plots of Mm Δ*facl6* vs Mm wt infected cells showing differential genes expression at 1, 6 and 24 hpi after performing DEGs analysis. Each dot represents one gene, plotted by log₂ fold change versus -log₁₀ P value. Selected cut-off for statistical significance (black dashed lines is abs(log2FC) > 0.5848 and padj < 0.05. (**B-E**) The profile of DEGs related to the ESCRT machinery (**B-C**) or the autophagy machinery (**D-E**) (previously defined gene lists) in infected cells compared to mock-treated cells was plotted as heatmap. B (**F**) The profile of DEGs related to the Esx-1 operon (previously defined gene lists) in Mm WT or Δ*facl6* during Dd infection time-course in Dd wt or *atg1*-KO cells, when compared to inocula, was plotted as heatmap. Blue: downregulated genes, and red: upregulated genes. (**G**) Schematic working model summarizing predicted Esx-1-associated secretion states based on transcriptional profiles. When Mm resides within an MCV, expression of the Esx-1 secretion system is detected, consistent with potential secretion of EsxA and its chaperone EsxB. In contrast, cytosolic bacteria show reduced or absent Esx-1 gene expression, suggesting a corresponding decrease in Esx-1-dependent secretion.

Given the central role of the damage-repair cycle in determining infection outcome, we next focused on host genes associated with ESCRT-mediated membrane repair and autophagy-related pathways. In Dd wt cells, ESCRT-associated genes showed a coordinated transcriptional upregulation as early as 1 hpi, followed by a return toward basal expression levels at later time points (**Fig. 3B**). This early response was markedly attenuated in the Dd *atg1*-KO background, consistent with altered damage handling in the absence of autophagy. Comparison of host transcriptional responses to Mm wt and Mm Δ*facl6* infection revealed largely overlapping expression patterns, apart from a distinct divergence at 6 hpi in both host backgrounds, as highlighted by LogFC cross-correlation analyses (**Fig. 3C**). A similar temporal pattern was observed for autophagy-associated genes, whose transcriptional dynamics closely mirrored those of ESCRT components (**Fig. 3D-E**). Together, these data indicate that infection with Mm Δ*facl6* is associated with altered timing and persistence of host damage-repair responses.

The early and sustained engagement of host membrane repair pathways prompted us to examine bacterial transcriptional programs associated with host membrane damage, focusing on the ESX-1 secretion system. Analysis of bacterial transcriptomes revealed that, already in broth-grown bacteria, most genes within the ESX-1 locus, including those encoding the membrane-damaging effector EsxA, were transcriptionally active (**Fig. S2C**). Comparison of Mm wt and Mm Δ*facl6* inocula further showed that the majority of ESX-1-associated genes were transcriptionally upregulated in the absence of *facl6*, except for the *espACD* operon, which was downregulated (**Fig. S2D**). During infection of Dd wt cells, Mm wt strongly induced the entire ESX-1 locus, including the associated operon *espACD*, and this induction was sustained until 24 hpi (**Fig. 3F**). In Dd *atg1*-KO cells, Mm wt also showed early ESX-1 induction at 1 hpi, but this response was transient and markedly reduced at later time points. In contrast, Mm Δ*facl6* displayed a profoundly altered ESX-1 transcriptional profile in both host backgrounds, characterized by globally attenuated induction of the ESX-1 locus throughout infection, with the notable exception of the *espACD* operon during infection of Dd wt cells. These patterns indicate that absence of FACL6 is associated with dysregulated ESX-1 transcription. Together with the compartmentalization phenotypes observed in wt and *atg1*-KO Dd cells, these transcriptional patterns are consistent with the working model proposed in **Fig. 3G**, in which ESX-1 activity is selectively engaged during the intravacuolar stage of infection and becomes dispensable following bacterial access to the cytosol.

### Mm Δ*facl6* induces enhanced and sustained membrane damage during infection

Based on the compartment- and time-dependent transcriptional signatures observed by dual RNA-seq, we next directly assessed membrane damage and cytosolic exposure during infection with Mm wt, Mm Δ*facl6*, Mm ΔRD1 and the complemented Mm Δ*facl6::facl6* strain.

To this end, we first monitored the recruitment of host damage and repair markers to the MCV (**Fig.4A-F; Fig S3**). In Dd, recruitment of the ESCRT-associated protein Alix occurs early at sites of membrane damage, whereas polymerization of the ESCRT-III component Chmp4/Vps32 marks later stages of membrane repair (López-Jiménez et al., 2018, Raykov et al., 2023). Quantitative imaging revealed a significantly higher proportion of Alix-GFP-positive MCVs in cells infected with Mm Δ*facl6* compared to Mm wt throughout the infection time-course. Alix recruitment during infection with the complemented Mm Δ*facl6::facl6* strain remained significantly elevated relative to Mm wt (**Fig. 4A-B; Fig. S3A**). In contrast, recruitment of Chmp4/Vps32-GFP was increased during Mm Δ*facl6* infection but restored to Mm wt levels in the complemented strain (**Fig. 4C-D; Fig. S3B**). Recruitment of both reporters was strongly reduced during infection with the ESX-1 deficient Mm ΔRD1 mutant. In agreement with these observations, analysis of infected iBMDMs revealed increased recruitment of Galectin-3 (Gal-3), a cytosolic lectin that recognizes luminal glycans exposed upon phagosomal membrane damage. The Gal-3 signal was significantly elevated on MCVs containing Mm Δ*facl6*, whereas it was rarely detected on compartments containing Mm ΔRD1 (**Fig. S4A-B**). Gal-3 recruitment was most pronounced at early stages of infection, mirroring the temporal dynamics observed for ESCRT reporters recruitment in Dd and supporting enhanced early membrane damage during Mm Δ*facl6* infection.

**Figure 4.**
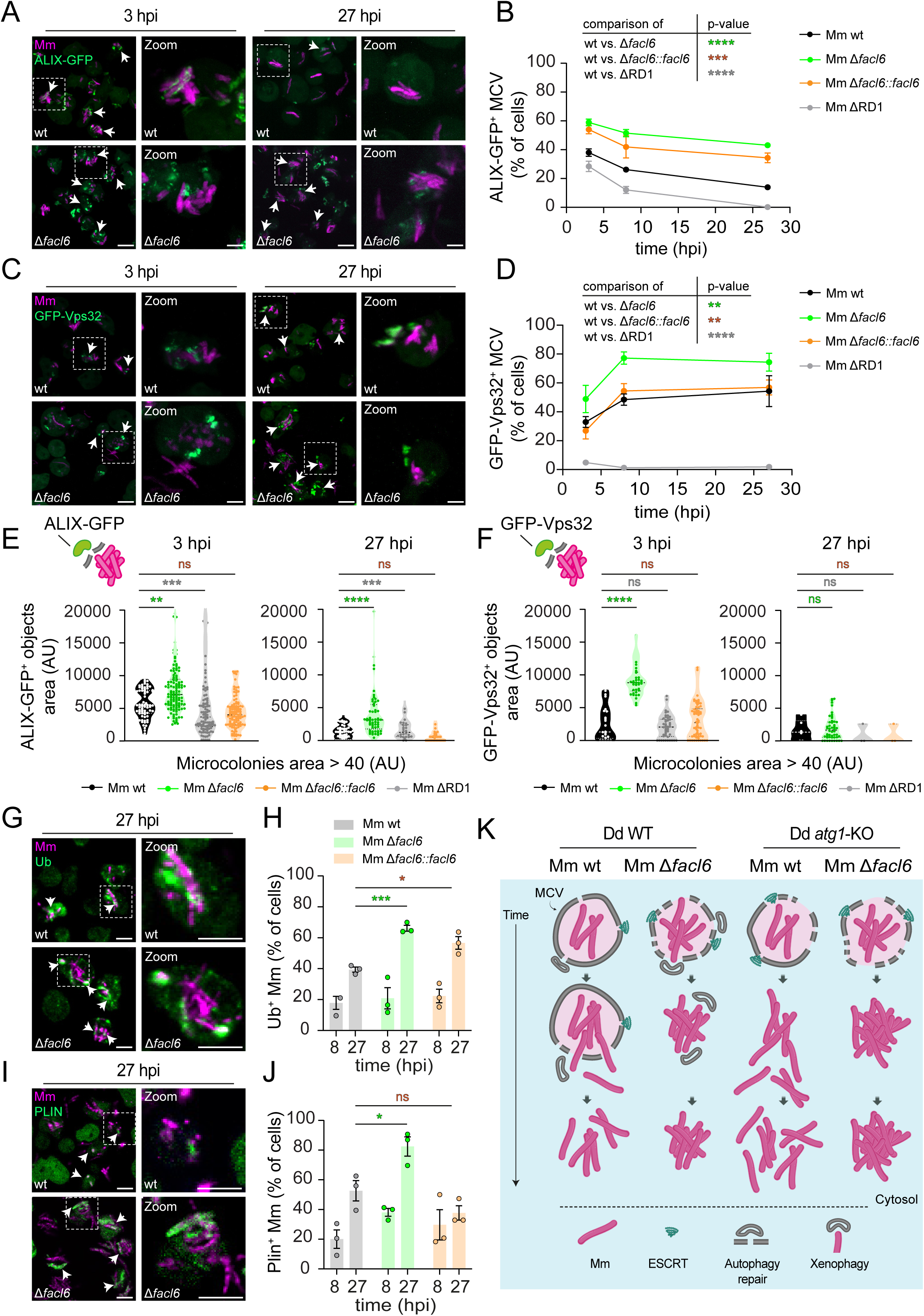
Mm Δ*facl6* induces exacerbated damage to its MCV, escaping earlier to the cytosol. **(A)** Live imaging of Alix-GFP recruitment to intracellular Mm in Dd. Arrows indicate Alix-GFP– positive puncta co-localizing with bacteria. (**B**) Quantification of Alix-GFP–positive MCVs. (**C**) Live imaging of GFP-Vps32 recruitment to intracellular Mm. Arrows indicate GFP-Vps32–positive co-localizing with bacteria. (**D**) Quantification of GFP-Vps32–positive MCVs. (**E**) Correlation between the number of Alix-GFP–positive puncta per cell and intracellular microcolonies larger than 40 AU. (F) Correlation between the number of GFP-Vps32–positive puncta per cell and intracellular microcolonies larger than 40 AU. (**G**) Immunofluorescence images of Ubiquitin (Ub) recruitment to intracellular Mm. Arrows indicate Ub-positive bacteria. (H) Quantification of Ub-positive bacteria. (**I**) Images of RFP-Plin recruitment to intracellular Mm. Arrows indicate Plin-positive bacteria. (**J**) Quantification of Plin-positive bacteria. (**K**) Working model of the different compartments and the differential recruitment of host-defense machineries over the time-course of the infection depending on the host and the bacteria genetic background. Dd cells were infected with Mm wt, Δ*facl6*, Δ*facl6::facl6*, or ΔRD1 (BFP- or GFP-expressing as indicated). Live imaging and fixed-cell imaging were performed using a spinning disc microscope (scale bars, 5 µm). Quantifications were performed manually (B, D, H, J) or using CellPathFinder (Yokogawa) (E, F). Data represent mean ± SEM (N = 3) Statistical significance was determined using an unpaired two-tailed Student’s t-test (*p < 0.05, **p < 0.01; ***, p < 0.001, ****p < 0.0001, ns: not significant).

To assess whether the extent of membrane damage correlated with the microcolony phenotype observed for Mm Δ*facl6*, we next analyzed ESCRT recruitment as a function of bacterial cluster size and infection time. At early stages of infection (3 hpi), larger Mm Δ*facl6* clusters exhibited higher recruitment of both Alix-GFP and Chmp4/Vps32-GFP compared to size-matched clusters of Mm wt or Mm ΔRD1, reflecting enhanced early damage sensing and early engagement of membrane repair (**Fig. 4E-F**). At later stages of infection (27 hpi), Alix-GFP recruitment increased with bacterial cluster size for Mm wt, Mm Δ*facl6* and Mm ΔRD1. However, at comparable cluster sizes, Mm Δ*facl6* microcolonies consistently showed higher Alix association than microcolonies for Mm wt or Mm ΔRD1, throughout the infection time course (**Fig. 4E; Fig. S3C-D**). In contrast, Chmp4/Vps32-GFP recruitment at 27 hpi correlated with bacterial cluster size similarly across all strains, and the early difference observed for Mm Δ*facl6* was no longer apparent (**Fig. 4F; Fig. S3C-D**).

To further assess cytosolic exposure of intracellular bacteria, we next monitored recruitment of Ubiquitin (Ub) and Perilipin (Plin), two cytosolic markers associated with damaged MCV and cytosolic-exposed mycobacteria (Barisch et al., 2015b). Quantitative analysis revealed that Ub association with intracellular bacteria was comparable between Mm wt, Mm Δ*facl6*, and the complemented Mm Δ*facl6::facl6* strain at 8hpi. In contrast, at 27 hpi, a significantly higher proportion of Mm Δ*facl6* bacteria were ubiquitinated compared to Mm wt, with the complemented strain displaying an intermediate phenotype (**Fig. 4G-H**). Plin association with intracellular bacteria showed a trend toward increased recruitment to Mm Δ*facl6* at 8 hpi. At later time point, Mm Δ*facl6* exhibited significantly higher Plin recruitment compared to both Mm wt and the complemented strain (**Fig. 4I-J**). Similar patterns were observed in iBMDMs, where staining for Ub and PLIN2 likewise revealed enhanced association with Mm Δ*facl6* and minimal recruitment to Mm ΔRD1 (**Fig. S4C-F**). Finally, consistent with increased cytosolic exposure, LC3 staining in iBMDMs demonstrated that a substantial fraction of Mm Δ*facl6* bacteria was associated with LC3-positive membranes, indicative of autophagy-related host responses, whereas Mm ΔRD1 showed limited LC3 recruitment (**Fig. S4G-H**).

Because early cytosolic access and extensive membrane damage have been associated with compromised host cell integrity in mammalian cells (Beckwith et al., 2020), we next assessed whether absence of FACL6 also impacted plasma membrane integrity in infected macrophages. iBMDMs infected with Mm wt, Mm Δ*facl6*, or Mm ΔRD1 were stained with PyroMarker, which selectively labels cells with compromised plasma membrane integrity. Quantitative analysis revealed a significantly higher proportion of plasma membrane–damaged cells during infection with Mm Δ*facl6*, whereas Mm ΔRD1 and the complemented Mm Δ*facl6::facl6* strain showed levels similar to wt (**Fig. S5A-B**). In addition, morphological analysis further suggested a trend toward increased features associated with pyroptotic and necrotic cell death during Mm Δ*facl6* infection (**Fig. S5C**), accompanied by a corresponding, though not statistically significant, increase in the overall number of dead cells compared to Mm wt and Mm ΔRD1 infections (**Fig. S5D**).

Together, these results demonstrate that during intracellular infection, absence of FACL6 leads to enhanced and sustained MCV membrane damage, resulting in premature and increased bacterial exposure to the cytosol, engagement of host cytosolic defense pathways as well as increased plasma membrane disruption. Based on the combined bacterial and host transcriptomic profiles, together with the compartment-specific infection phenotypes, these observations are integrated in the infection time-course working model presented in **Fig. 4K**. In this model, Mm Δ*facl6* is associated with earlier engagement of host damage–repair pathways and earlier access to the cytosol in Dd wt cells, where bacteria persist as intracellular microcolonies. In the Dd *atg1*-KO background, Mm Δ*facl6* similarly gains cytosolic access but, despite the absence of xenophagic restriction, grows relatively slowly and predominantly as microcolonies.

### Absence of FACL6 perturbs the mycobacterial cell envelope and increases drug susceptibility in axenic culture

The cytosolic microcolony phenotype of Mm Δ*facl6* raised the question why the mutant fails to proliferate maximally even after gaining access to the cytosol, a compartment that supports unrestricted growth of Mm wt in Dd *atg1*-KO cells. This limitation could reflect increased susceptibility to host cytosolic antibacterial defenses and/or an intrinsic constraint on bacterial growth, such as altered bacterial physiology or metabolism. Notably, bacterial surface and envelope properties are key determinants of how cells interact and organize; in other bacteria, variation in surface adhesion factors influences microcolony architecture, and in mycobacteria, alterations in cell-envelope polysaccharides change surface properties and intercellular interactions (Lian et al., 2025, Duvernoy et al., 2018). To distinguish between host-driven and intrinsic bacterial contributions to the Mm Δ*facl6* growth defect, we examined the mutant under axenic growth conditions in 7H9/OADC medium. Scanning electron microscopy (SEM) revealed pronounced surface abnormalities in Mm Δ*facl6*. While Mm wt bacteria displayed a characteristically smooth and continuous outer surface, Mm Δ*facl6* exhibited a markedly rough appearance with irregular surface protrusions and localized bulges (**Fig. 5A**). To characterize molecular signatures associated with this altered envelope phenotype, we compared at global level the transcriptional profiles of Mm Δ*facl6* and Mm wt grown in broth. This analysis revealed that the most strongly differentially expressed genes, both up- and downregulated, were enriched for PE/PPE family proteins and cell wall–associated functions (**Fig. S6A-B**). Notably, approximately half of the upregulated genes and nearly 60% of the downregulated genes mapped to these two functional categories, prompting a focused analysis of cell wall-related transcriptional changes. Consistent with this enrichment, Mm Δ*facl6* displayed a distinct transcriptional profile of cell wall-associated genes, with specific up- and downregulated factors linked to cell-envelope biogenesis and remodeling (**Fig. 5B**). Among the most strongly upregulated genes was *pks13*, which encodes a key polyketide synthase required for mycolic acid biosynthesis, suggesting compensatory induction of mycolic acid production in response to envelope perturbation. In addition, several differentially expressed genes mapped to core peptidoglycan pathways, including *murA*, *murI*, and *murT*, consistent with broader remodeling of the bacterial cell wall in the absence of FACL6. We next assessed whether these structural and transcriptional alterations translated into functional changes in envelope integrity using an antibiotic susceptibility assay under axenic conditions. As controls, we included Mm Δ*tesA*, a mutant with established cell-envelope defects linked to impaired lipid biogenesis (Chavadi et al., 2011, Alibaud et al., 2011), and Mm ΔRD1, which also exhibits altered envelope-associated properties (Abdallah et al., 2019). Across all compounds tested, including isoniazid, vancomycin, penicillin/streptomycin, and moxifloxacin, Mm *Δfacl6* displayed increased susceptibility compared to Mm wt, similarly to the control mutants (**Fig. 5C**). These results are consistent with compromised cell-envelope integrity and increased permeability.

**Figure 5.**
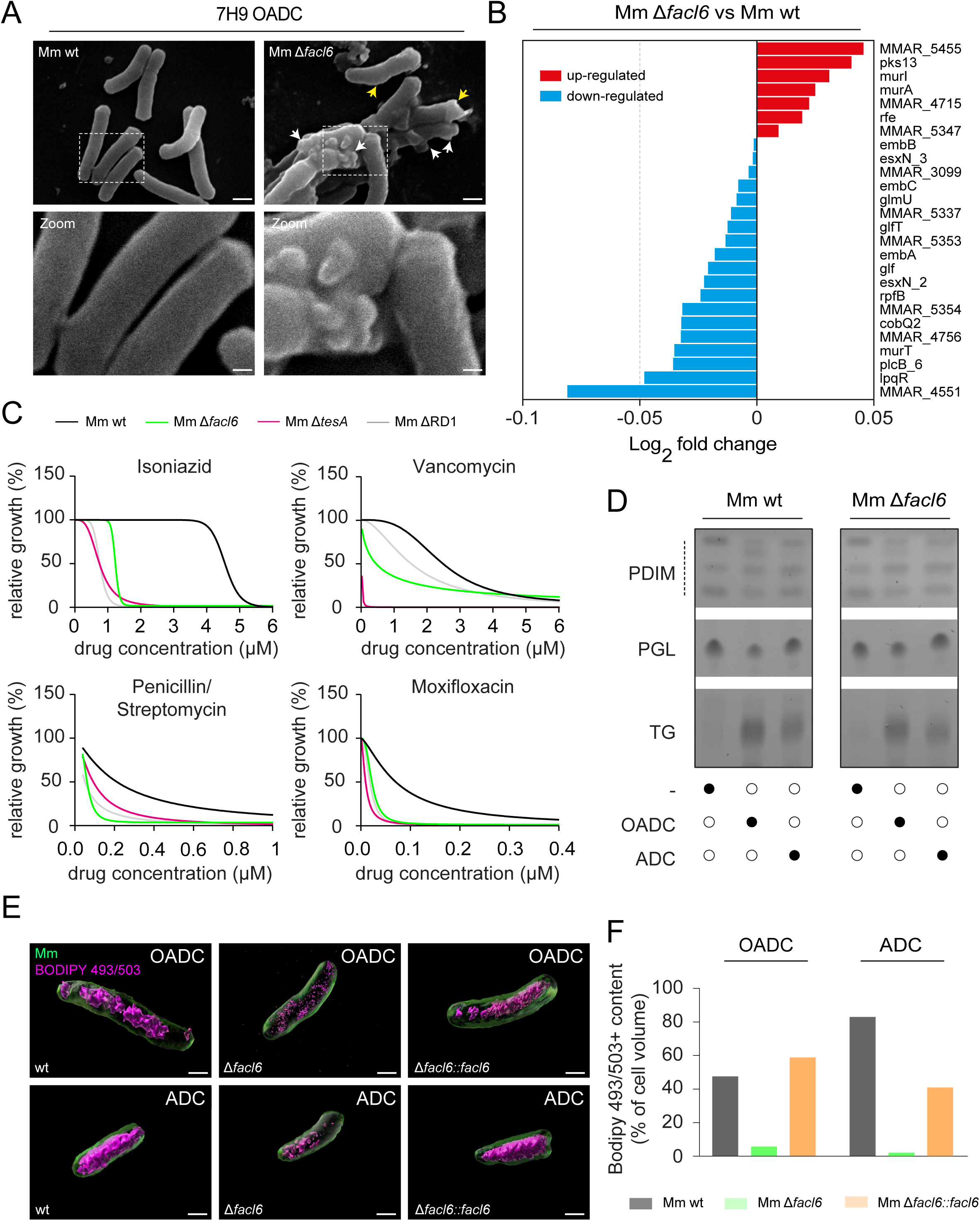
FACL6 deletion correlates with alteration in Mm cell wall and neutral lipid storage. **(A)** Scanning electron microscopy (SEM) of Mm wt or Mm Δ*facl6* grown in axenic medium (7H9/OADC). Yellow arrows indicate irregular and rough regions on the bacterial surface, while white arrows highlight vesicular budding (scale bars, 1 µm; zoom, 0.1 µm). (**B**) Cell wall related genes (Uniprot “cell wall” gene list) differentially expressed between Mm wt or Mm Δ*facl6* grown in axenic medium. Blue: upregulated in Mm Δ*facl6*, red: downregulated in Mm Δ*facl6* compared to Mm wt. (**C**) Luminescent Mm wt, Δ*facl6*, Δ*tesA* or ΔRD1 were assessed for their *in vitro* susceptibility to Isoniazid, Vancomycin, Penicillin/Streptomycin or Moxifloxacin. Bacterial luminescence was measured using a microplate reader. Relative growth was calculated by normalizing luminescence values to untreated controls (100% growth) and to samples treated with 2 µM Rifampicin (0% growth). Dose–response curves illustrate relative bacterial growth plotted against antibiotic concentration. One representative experiment of two independent experiments is shown. (**D**) TLC of total lipids extracted from Mm wt and Δfacl6 grown in different culture media . (7H12 containing lipid-rich (OADC), lipid-depleted (ADC) or no enrichment (-))Lipid species including PDIM, PGL and TG were resolved and visualized.(**E**) Visualization and (**F**) quantification of bacterial ILI *in vitro* under different nutrient conditions. GFP-expressing Mm wt and Δ*facl6* were cultured in 7H9 medium supplemented with OADC or ADC for 24 hours. Bacteria were stained with Bodipy 493/503 and fixed.Images were acquired using a spinning disc confocal microscope operated in super-resolution by optical re-assignment (SoRa) mode. Representative 3D images were generated with Imaris. (**F**) Quantification of 3D image analysis. Volumetric ILIs content was quantified with Imaris. Data shown are from one representative experiment.

To determine whether major outer-envelope lipids were affected by *facl6* deletion, we compared PDIM and PGL profiles by thin-layer chromatography (TLC) in Mm wt and Mm Δ*facl6* grown under lipid-rich (OADC), lipid-depleted (ADC), or minimal conditions. No major differences were detected between the two strains under any of these conditions, indicating that the observed phenotypes are not driven by gross perturbations of these outer-envelope lipids (**Fig. 5D; Fig. S6C-D**). As expected, TG abundance in Mm wt varied with nutrient availability, with highest levels in OADC, reduced levels in ADC, and minimal levels under minimal conditions. Across all growth conditions, however, TG levels appeared modestly reduced in Mm Δ*facl6* compared to Mm wt, compatible with the proposed function of FACL6 in coupling fatty acid import and CoA ligation that fuels intracellular lipid inclusions (ILI) genesis (**Fig. 1A**). To further characterize neutral lipid storage at the single-bacterium level, we next quantified ILI in bacteria grown in axenic medium containing OADC or ADC and stained with the neutral lipid probe BODIPY 493/503, using super-resolution microscopy via optical re-assignment (SoRa) (**Fig. 5E**). Volumetric analysis revealed that Mm Δ*facl6* contained markedly fewer neutral lipids, presumably ILIs, than Mm wt in both growth conditions, as reflected by reduced BODIPY 493/503 positive content per bacterium (**Fig. 5F**). In contrast, the complemented Mm Δ*facl6::facl6* strain exhibited BODIPY 493/503 signal intensities comparable to Mm wt. Together, these results demonstrate that absence of FACL6 impairs neutral lipid accumulation independently of lipid availability in the growth medium.

Collectively, these findings show that absence of FACL6 causes intrinsic alterations in mycobacterial physiology, characterized by cell-envelope remodeling, increased permeability, and defective neutral lipid storage under axenic conditions. This raised the possibility that the mutant may be unable to appropriately utilize host-derived lipid substrates.

### Absence of FACL6 alters fatty acid-dependent growth and renders Mm sensitive to saturated fatty acids *in vitro*

To determine whether the intrinsic physiological alterations observed in Mm Δ*facl6* translate into altered growth behavior under defined nutrient conditions, we compared the growth kinetics of Mm wt and Mm Δ*facl6* in nutrient-rich versus minimal media. As expected, both strains displayed shorter generation times in rich conditions (ADC/OADC-supplemented 7H12 or 7H9) than in minimal media (**Fig. 6A-B**). Notably, under minimal conditions (7H12) the Mm Δ*facl6* mutant exhibited a significantly shorter generation time than Mm wt, indicating a relative growth advantage specifically under nutrient limitation and suggesting metabolic rewiring of the mutant. To directly test how specific host-relevant lipids influence growth, Mm wt, Mm Δ*facl6* and Mm Δ*facl6::facl6* were cultured in minimal 7H12 medium supplemented with increasing concentrations of cholesterol (100, 200, or 1000 μM), oleate (10, 20, or 100 μM), or palmitate (10, 20, or 100 μM). Growth curves are shown in **Fig. S7**, and corresponding AUC values are summarized in **Fig. 6C-E**. Cholesterol supplementation had only minor effects on growth of all strains across concentrations tested, consistent with sterols supporting only moderate growth under these conditions (**Fig. 6C**). In contrast, supplementation with fatty acids significantly enhanced growth of Mm wt and Mm Δ*facl6* and Mm Δ*facl6::facl6* (**Fig. 6D,E**). At intermediate concentrations of both oleate and palmitate, Mm Δ*facl6* grew significantly better than Mm wt. However, this growth advantage was lost at high concentrations of palmitate. Exposure to the highest concentration of palmitate resulted in a pronounced growth inhibition that was specific to the Mm Δ*facl6* mutant and only partially alleviated in the complemented strain (**Fig. 6E**). This inhibitory effect was not observed at the highest oleate concentration (**Fig. 6D**), nor in Mm wt, indicating a selective sensitivity of Mm Δ*facl6* to saturated fatty acids. To determine whether palmitate-induced growth inhibition could be mitigated by alternative lipid substrates, bacteria were cultured in 7H12 medium containing increasing concentrations of palmitate together with either cholesterol or oleate (**Fig. 6F-G**). As observed previously, Mm Δ*facl6* displayed concentration-dependent growth inhibition in the presence of palmitate. Addition of cholesterol led to a modest improvement in growth relative to palmitate alone but did not restore growth to levels observed in the absence of palmitate (**Fig. 6F**). In contrast, co-supplementation with oleate significantly improved growth of Mm Δ*facl6* in the presence of palmitate (**Fig. 6G**). Together, these results show that saturated fatty acid toxicity associated with absence of FACL6 can be effectively counterbalanced by a monounsaturated fatty acid, highlighting a key role for FACL6 in maintaining lipid metabolic balance under conditions of fatty acid excess (**Fig. 6H**).

**Figure 6.**
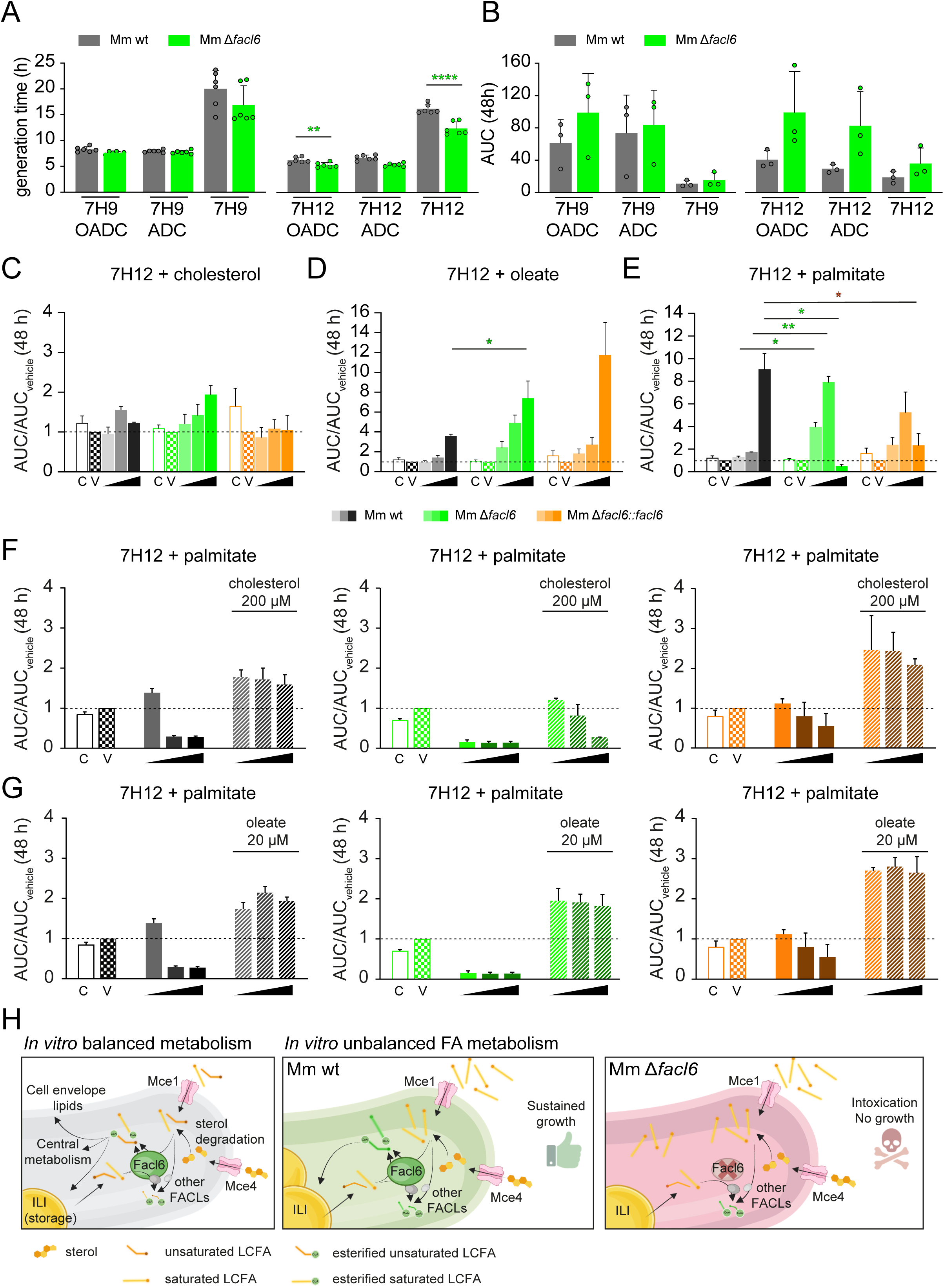
FACL6 deletion alters Mm lipid turnover and sensitivity to host-relevant fatty acids. **(A)** Generation time of GFP-expressing Mm wt or Δ*facl6* were grown in 7H9 or 7H12 containing lipid-rich (OADC), lipid-depleted (ADC) or no enrichment. (**B**) Area under the curves (AUC) of the corresponding growth curves. (**C**-**E**) Growth of GFP-expressing Mm wt, Δ*facl6*, or Δ*facl6::facl6* in 7H12 medium alone (control: C) or supplemented with increasing concentrations of cholesterol (**C**), oleate (**D**), or palmitate (**E**), or with the corresponding vehicle control (vehicle: V). (**F to G**) Growth of GFP-expressing Mm wt, Δ*facl6*, or Δ*facl6::facl6* in 7H12 medium containing increasing concentrations of palmitate in the absence or presence of additional cholesterol (**F**) or oleate (**G**). Fluorescence signal was measured using a microplate reader. Generation times were estimated over 72 hours growth, with each dot being an individual replicate (**A**). AUCs of growth curves over 48 h were plotted. Shown are mean ± SEM, N=3. Statistical significance was determined using an unpaired two-tailed Student’s t-test, (*p < 0.05, **p < 0.01). (**H**) Schematic representations of the working model related to *in vitro* balanced versus unbalanced saturated fatty acid metabolism in Mm.

### Mm Δ*facl6* infection induces a distinct host lipid metabolic transcriptional response

The *in vitro* growth phenotypes observed for Mm Δ*facl6* suggested that absence of FACL6 profoundly alters bacterial metabolism, raising the question of whether these metabolic differences are also reflected in the host cell environment during infection. To address this, we analyzed the host-side component of our dual RNA-seq dataset, focusing on transcriptional responses of Dd cells infected with Mm wt or Mm Δ*facl6* (**Fig. 7**). Pathway enrichment analysis was performed by comparing infected cells to mock-treated controls at 6 hpi, the time point previously identified as critical for divergence between infection outcomes (**Fig. 3A**). Several pathways were similarly enriched in Dd cells infected with Mm wt or Mm Δ*facl6*, including endocytosis, phagosome maturation, autophagy-related pathways, and pathways associated with membrane remodeling and intracellular trafficking (**Fig. 7A-B**). In addition to these shared responses, Dd cells infected with Mm Δ*facl6* showed specific enrichment of pathways linked to lipid metabolism, including α-linolenic acid metabolism, pantothenate and CoA biosynthesis, and biosynthesis of unsaturated fatty acids (**Fig. 7B**).

**Figure 7.**
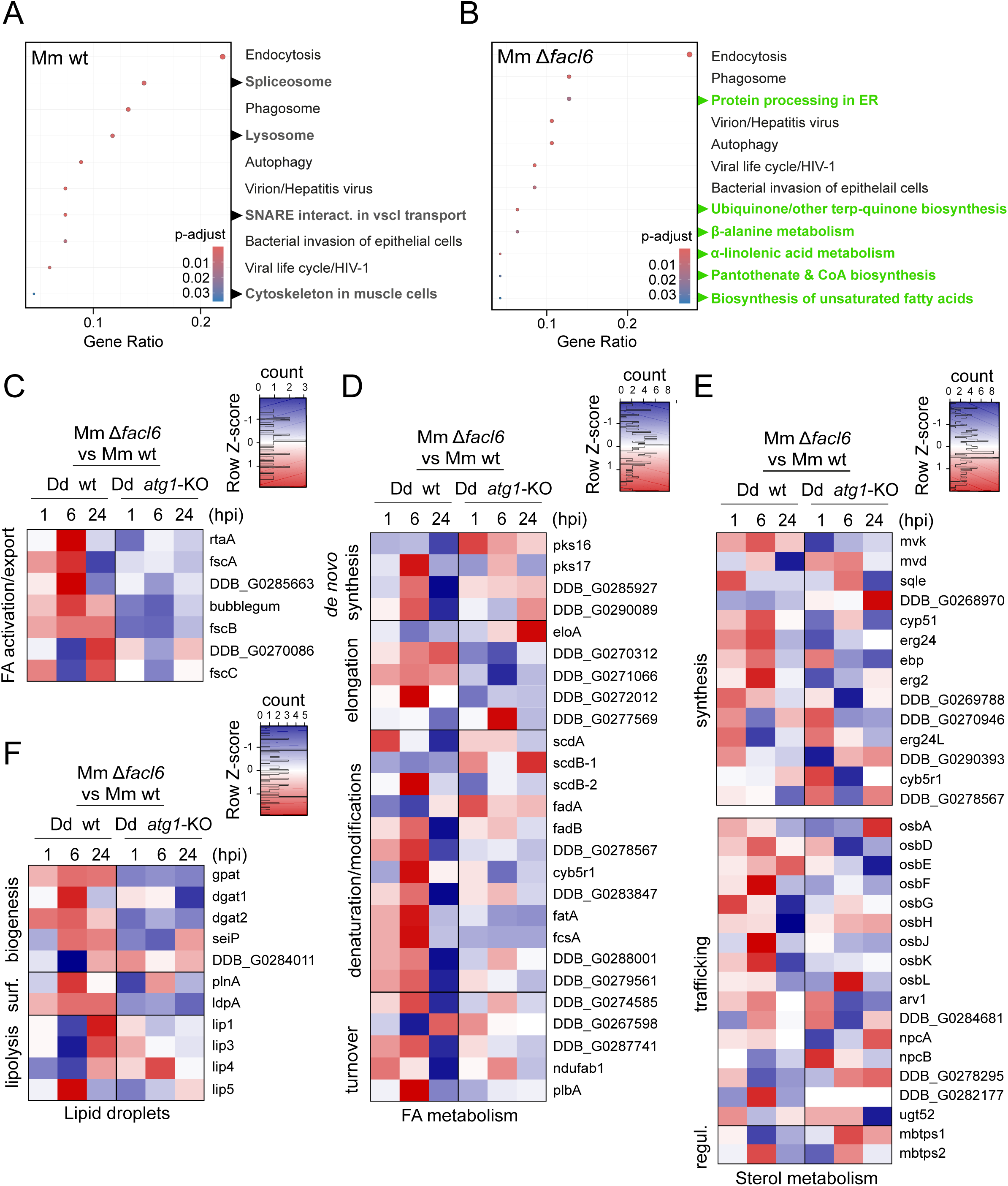
Dd cells infected with Mm. Δ***facl6* exhibit a rewiring of their lipid metabolism program at the transcriptomic level.** (**A-B**) Gene set enrichment analysis (GSEA) was performed with the Kyoto Encyclopedia of Genes and Genomes (KEGG) annotation, significantly enriched terms (*p < 0.05) at 6 hpi for both Mm wt (**A**) or Mm Δ*facl6* (**B**) infected cells when compared to mock. Results are shown as dotplots. The color code of the dots stands for p-adjust values. Bold black: pathways specifically enriched in Mm wt-infected conditions, bold green: pathways specifically enriched in Mm Δ*facl6*-infected conditions. The profile of DEGs related to fatty acid activation (**C**) or metabolism (**D**), as well as sterol metabolism (**E**) and LDs (**F**) (previously defined gene lists) in Mm Δ*facl6* vs Mm wt infected cells was plotted as heatmap. Blue: downregulated genes, red: upregulated genes.

Prompted by this pathway enrichment, we thus examined curated gene sets associated with host lipid metabolism. Analysis of genes involved in fatty acid activation, transport and metabolism revealed a broad transcriptional upregulation in Dd wt cells infected with Mm Δ*facl6*, which was already detectable at 1 hpi and became most pronounced at 6 hpi (**Fig. 7C-D**). This response was largely absent in the Dd *atg1*-KO background, where bacteria rapidly access the cytosol and infection outcomes are less divergent from Mm wt, indicating that the metabolic host response is tightly linked to the infection status. A similar pattern was observed for genes involved in sterol metabolism (**Fig. 7E**). We next examined genes associated with LD biology. As observed for fatty acid and sterol metabolism, most LD-associated genes were upregulated at 1 and 6 hpi in Dd wt cells infected with Mm Δ*facl6* compared to Mm wt (**Fig. 7F**). In contrast, genes linked to lipolysis showed a distinct temporal pattern: most lipases-encoding genes were downregulated at early time points and became upregulated at 24 hpi. Notably, *lip5* represented an exception, displaying increased expression at 6 hpi followed by downregulation at later stages of infection (**Fig. 7F**). Together, these results show that Mm Δ*facl6* infection is associated with enhanced activation of host lipid metabolic pathways at an early stage of infection, likely defining a host metabolic environment distinct from that encountered during Mm wt infection.

### The Mm transcriptomic profile highlights compartment-specific lipid metabolic programs in intracellular mycobacteria

To determine whether host lipid metabolic transcriptional responses observed during Mm Δ*facl6* infection are associated with distinct bacterial transcriptional programs, we next analyzed the mycobacterial component of the dual RNA-seq dataset (**Fig. 8**). We first performed principal component analysis (PCA) on the Mm transcriptomes collected during infection of Dd wt and Dd *atg1*-KO cells. PCA revealed a clear stratification of samples according to bacterial genotype, infection time point, and host genetic background, with Mm wt and Mm Δ*facl6* forming two distinct clusters (**Fig. 8A**). This segregation indicates that mycobacterial transcriptional responses are influenced by both intracellular compartment and infection stage.

**Figure 8.**
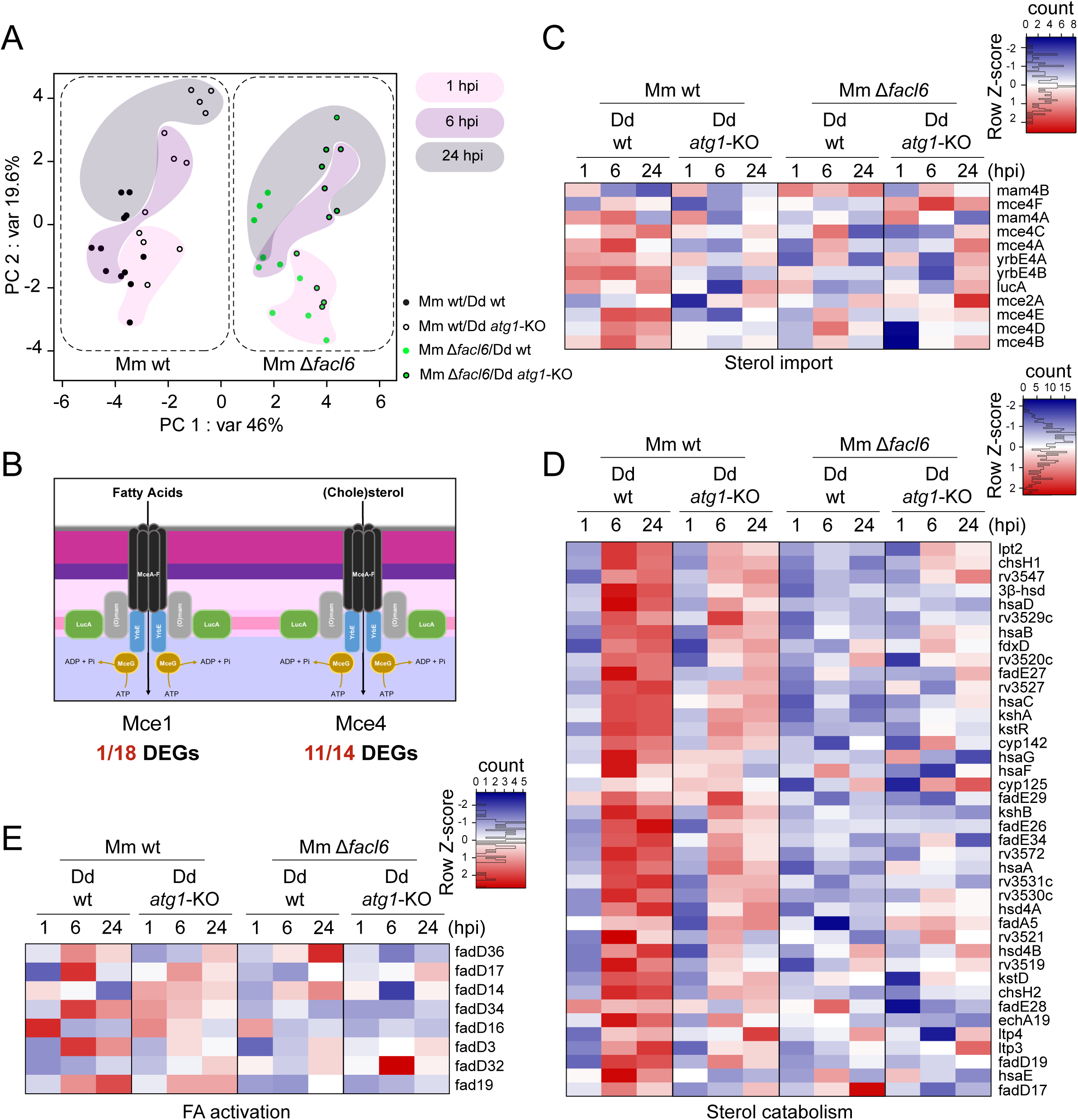
Mm Δ*facl6* specific transcriptomic signature highlights stage-specific and compartment-specific lipid metabolic program by intracellular mycobacteria. (**A**) Principal Component Analysis (PCA) show strain and time-point specific stratifications when principal component 1 (representing 46% of the total variance) and principal component 2 (representing 19% of the total variance) are plotted. (**B**) Schematic representation of Mm protein complexes dedicated to fatty acids (Mce1) and sterol (Mce4) import, and the number of DEGs in Mm Δ*facl6* vs wt over the total number of genes required for the synthesis of these systems (18 among the *mce1* operon, 14 among the *mce4* operon) during the whole Dd infection time-course. The profile of DEGs related to sterol import (**C**) and catabolism (**D**) or fatty acid activation (FAAL/FACL) (**E**) (previously defined gene lists) in Mm wt or Δ*facl6* during Dd infection time-course in wt or *atg1*-KO cells, when compared to inocula, was plotted as heatmap. Blue: downregulated genes, red: upregulated genes.

Given the *in vitro* fatty acid sensitivity of Mm Δ*facl6* and the host lipid metabolic signatures identified in **Fig. 7**, we next focused on bacterial pathways involved in host-derived lipid acquisition. Mm encodes two major lipid uptake systems: Mce1, primarily implicated in fatty acid uptake, and Mce4, which mediates sterol import (**Fig. 8B**). Analysis of differentially expressed genes (DEG) across all infection conditions revealed a striking disparity between these systems. Whereas no core components of the Mce1 transporter were transcriptionally regulated, most genes encoding components of the Mce4 system were differentially expressed during infection (**Fig. 8B**). The only DEG functionally linked to fatty acid uptake was *lucA*, a gene encoding a subunit known to interface with both Mce1- and Mce4-dependent lipid transport (Nazarova et al., 2017). We therefore examined sterol import and catabolism in greater detail. In Mm wt infecting Dd wt cells, genes encoding the Mce4 sterol uptake system as well as the full sterol catabolic program were strongly induced from 6 hpi to 24 hpi (**Fig. 8C-D**). This time window corresponds to the intravacuolar phase of infection, consistent with active sterol acquisition and utilization within the MCV. In contrast, during infection of Dd *atg1*-KO cells, where bacteria prematurely access the cytosol, the induction of sterol-related genes in Mm wt was detectable but markedly reduced. In the Mm Δ*facl6* mutant, sterol-associated transcriptional responses were strongly attenuated. In both Dd wt and Dd *atg1*-KO backgrounds, Mm Δ*facl6* showed little to no induction of genes involved in sterol import or degradation throughout the infection time course (**Fig. 8C-D**). This contrasts with the robust sterol-related transcriptional response observed in Mm wt during the intravacuolar phase of infection and indicates that sterol-associated gene induction correlates with the intravacuolar micro-environment.

We next examined transcriptional regulation of genes involved in fatty acid activation, focusing on the FAAL/FACL family. Although Mm encodes 33 FAAL/FACL enzymes, only 9 were differentially expressed under the infection conditions analyzed (**Fig. 8E**). Like the sterol-related genes, FAAL/FACL genes were upregulated in Mm wt during infection of Dd wt cells, showed weaker induction in Dd atg1-KO cells, and exhibited minimal transcriptional regulation in the Mm Δ*facl6* mutant in both host backgrounds (**Fig. 8E**).

Together, these results indicate that lipid-associated transcriptional responses in intracellular Mm are strongly dependent on compartment and infection stage. In Mm wt, sterol import and catabolic genes are preferentially induced during the intravacuolar phase, whereas cytosolic localization is associated with attenuated induction. In the absence of FACL6, these compartment-associated transcriptional patterns are altered, most likely as a consequence of the premature MCV damage and escape to the cytosol.

### Intracellular compartmentalization shapes lipid acquisition and neutral lipid storage by mycobacteria

Based on our transcriptomic and physiological analyses, we hypothesized that lipid acquisition and storage by intracellular mycobacteria are tightly constrained by intracellular compartmentalization. Specifically, our data predict that the intravacuolar environment correlates with neutral lipid accumulation (**Fig. 8D**), whereas cytosolic exposure is associated with reduced lipid storage and increased sensitivity to lipid overload. We therefore examined how compartmentalization impacts lipid acquisition and ILI during infection. Transmission electron microscopy (TEM) of infected Dd cells at 24 hpi revealed marked differences in ILI morphology. Whereas Mm wt bacteria contained well-defined, compact ILI, Mm Δ*facl6* bacteria displayed more fragmented lipid inclusions (**Fig. 9A**). These observations indicate that although ILI formation appears reduced in the Mm Δ*facl6* mutant under lipid-rich axenic conditions (**Fig. 5E-F**), it can form ILI during intracellular infection, albeit with altered structural organization. This raised the question of whether Mm Δ*facl6* differs from Mm wt in its ability to acquire host-derived fatty acids and channel them into neutral lipid storage during infection. In this context, we employed a live imaging approach using BODIPY-C12-labeled host lipids. Dd cells expressing AmtA-GFP were preloaded with palmitate and BODIPY-C12 (i.e. a fluorescent fatty acid analogue) prior to infection with BFP-expressing Mm wt, Mm Δ*facl6*, or the complemented Mm Δ*facl6::facl6* strain, and infections were followed over time (**Fig. 9B-C**). Incubation of Dd with palmitate supports the incorporation of BodipyC12 into TGs and LDs, which helps tracking the lipid flow from the host to the pathogen (Barisch et al., 2015b). AmtA-GFP marks endosomal membranes in Dd and was used to delineate the MCV (**Fig. 9B**; (Barisch et al., 2015b)). Quantitative analysis revealed that at all time points examined, a significantly lower proportion of BODIPY-C12-positive Mm Δ*facl6* bacteria was observed compared to Mm wt, with the complemented strain showing intermediate to wt-like levels (**Fig. 9D**). For both Mm wt and Mm Δ*facl6* strains, the overall proportion of BODIPY-C12-positive bacteria decreased over time, consistent with turnover or redistribution of the fluorescent lipid tracer during infection. Regarding their intracellular localization, BODIPY-C12-positive Mm wt and Mm Δ*facl6* bacteria were detected in similar proportions within vacuolar and cytosolic compartments at early time points. At later stages of infection, however, the fraction of BODIPY-C12-positive Mm wt bacteria were predominantly cytosolic, whereas it remained more evenly distributed between vacuolar and cytosolic locations for Mm Δ*facl6* bacteria (**Fig. 9E**). To assess neutral lipid storage independently of exogenous lipid labeling, we also performed SBF-SEM of intracellular bacteria. Volumetric reconstruction of infected cells suggested that Mm Δ*facl6* bacteria tended to contain less total ILI volume than Mm wt during intracellular infection (**Fig. 9F-H**). Interestingly, comparison of Mm wt bacteria residing in different compartments indicated a possible reduction in ILI-associated volume in cytosolic compared to vacuolar Mm wt . Together, these observations support the idea that cytosolic localization is associated with a marked reduction in neutral lipid storage, consistent with a shift away from lipid accumulation once bacteria exit the MCV.

**Figure 9.**
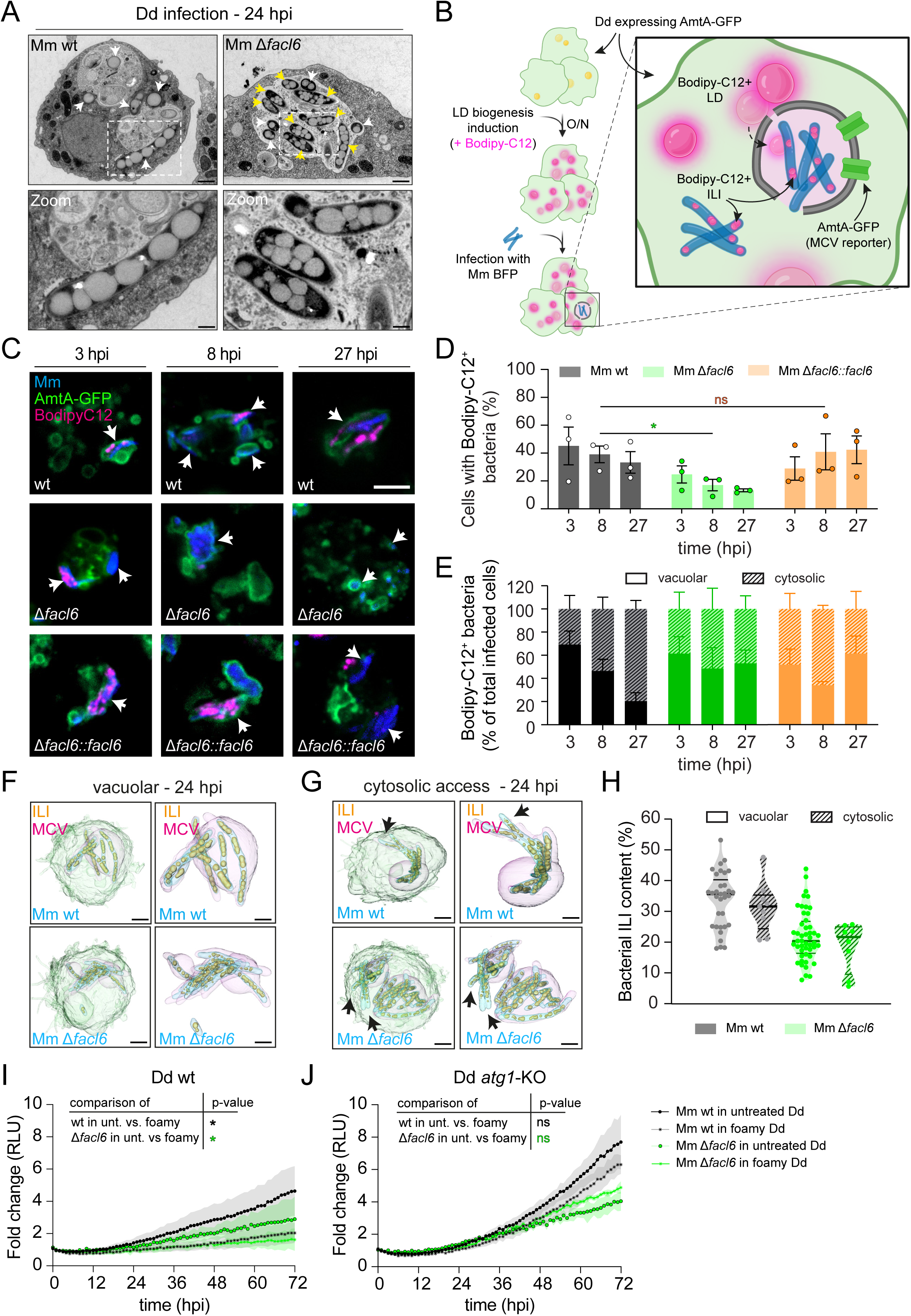
Mm host-derived lipid acquisition is compartment-dependent. (**A**) Transmission electron microscopy (TEM) images of Dd infected cells with Mm wt or Mm Δ*facl6* fixed at 24 hpi (scale bars, 3 µm (inset = 1 µm). White arrows point to Mm within infected cells. Yellow arrows show fragmented ILIs in Mm Δ*facl6*. (**B**) Schematic representation of the experimental setup for panels C to E. (**C**) Live imaging of Dd infected with Mm wt, Δ*facl6*, or Δ*facl6::facl6* at 3, 8, and 27 hpi. Dd expressing AmtA-GFP was preincubated with Bodipy-C12 prior to infection. Samples were imaged using a spinning disc microscope (scale bars, 5 µm). (**D-E**) Quantification of the proportions of cells with Bodipy-C12 positive bacteria (D) or the proportions of Bodipy-C12 positive bacteria located either within or outside the AmtA-GFP positive vacuole (E). (**F-G**) 3D-reconstruction of bacterial ILI in Dd using SBF-SEM. Cells were infected with Mm wt and Δ*facl6*, respectively, fixed, and imaged at 24 hpi. (**H**) Volumetric quantification of lipid content normalized on bacteria volume in vacuolar and cytosolic host cell compartment. Data from two independent biological experiments were pooled. (**I**-**J**) Intracellular growth of Mm wt and Δ*facl6* in Dd wt or *atg1*-KO with or without a foamy phenotype. To induce an excess of neutral lipid accumulation in infected cells, Dd cells were pre- incubated with exogenous palmitate (in excess) for 4 hours prior to the infection and maintained all along the time-course of the infection. Dd wt (I) or *atg1*-KO (J) cells were then infected with luminescent Mm wt and Δ*facl6*. Luminescence was measured for 72 hours using a microplate reader. Shown are mean fold changes ± SEM, N=3. Statistical significance was calculated using a two-way ANOVA with Fisher’s LSD test (*p < 0.05, ns: not significant).

Finally, based on the compartment-specific differences in lipid acquisition and storage observed above, we predicted that excess host lipid availability would differentially impact intracellular bacterial growth depending on the balance between vacuolar and cytosolic phases of infection. To test this prediction, Dd wt or Dd *atg1*-KO cells were preloaded with palmitate to induce host neutral lipid accumulation prior to infection as in **Fig. 9B**, a foamy phenotype artificially maintained throughout the infection. In Dd wt cells, where bacteria experience prolonged intravacuolar phases across successive infection cycles, host foamy phenotype resulted in a pronounced intracellular growth defect for both Mm wt and Mm Δ*facl6* (**Fig. 9I**). In contrast, in the Dd *atg1*-KO background, which favors rapid and sustained cytosolic localization, the host foamy phenotype had a more limited impact on Mm wt growth and did not impair growth of Mm Δ*facl6* (**Fig. 9J**).

Together, these results validate a model in which bacteria intracellular compartmentalization governs host-pathogen lipid exchange and determines whether host lipid abundance supports bacterial growth or becomes detrimental. In this framework, FACL6 emerges as a key determinant enabling mycobacteria to tolerate and adapt to compartment-specific lipid conditions encountered during intracellular infection.

## Discussion

Intracellular mycobacteria are exposed to different metabolic environments depending on whether they reside within the MCV or gain access to the host cytosol. While host-derived lipids have long been recognized as important nutrients for intracellular mycobacteria (see for review (Wilburn et al., 2018, Laval et al., 2021)), how lipid availability and utilization are shaped by intracellular compartmentalization remains poorly understood. Here, we show that lipid metabolic programs in Mm are tightly linked to intracellular localization, with sterol acquisition and neutral lipid storage preferentially engaged during the intravacuolar phase of infection. Disruption of this coordination, as observed upon deletion of *facl6*, exacerbates the consequences of premature escape from the MCV.

### FACL6 supports envelope homeostasis and shapes virulence-associated phenotypes

Deletion of *facl6* resulted in pronounced alterations of the mycobacterial cell envelope. Under axenic conditions, Mm Δ*facl6* displayed marked ultrastructural abnormalities at the bacterial surface, accompanied by transcriptional remodeling of genes involved in mycolic acid synthesis and peptidoglycan biogenesis. These structural and transcriptional changes translated into functional defects, as evidenced by heightened susceptibility to multiple antibiotics, consistent with an increased envelope permeability (Chavadi et al., 2011, Alibaud et al., 2011). Although altered envelope-related phenotypes are often driven by gross loss of outer lipids (Jones et al., 2024, Lee et al., 2025, Weaver et al., 2025), PDIM/PGL profiles were not detectably altered, arguing instead for subtler remodeling of the mycomembrane rather than a catastrophic envelope failure. Consistent with this interpretation, deletion of *facl6* was associated with an altered differential expression of PE/PPE family genes, a hallmark transcriptional signature of envelope stress and surface remodeling in mycobacteria (Fishbein et al., 2015). PE/PPE proteins are enriched at the mycobacterial surface and have been implicated in modulating envelope permeability, host immune recognition, metabolism and virulence-associated secretion pathways (Fishbein et al., 2015, Ates, 2020, Boradia et al., 2025, Brennan, 2017, Block et al., 2026, Wang et al., 2020). Their coordinated upregulation in Mm Δ*facl6* further supports the idea that disruption of fatty acid activation perturbs envelope homeostasis and triggers compensatory surface remodeling programs.

In addition to these molecular and ultrastructural defects, loss of FACL6 profoundly altered the spatial organization of intracellular bacteria. Mm Δ*facl6* consistently formed compact, spherical microcolonies both in axenic culture and during infection, in contrast to the more aligned and elongated bacterial assemblies formed by Mm wt. Similar alterations in bacterial organization have been reported for Mtb mutants defective in mycolic acid modification or lipid metabolism, such as Δ*pcaA* or Δ*fadD26*, which form dense bacterial aggregates and display increased antibiotic susceptibility (Mishra et al., 2023). In that study, bacterial aggregates were shown to impose localized mechanical stress on host membranes and to elicit distinct host responses compared to more dispersed bacteria. The microcolony phenotype of Mm Δ*facl6*, together with its roughened surface architecture, could participate in amplifying focal membrane deformation and damage at the MCV interface, providing a physical complement to the biochemical envelope defects described above.

### Envelope perturbation is associated with altered ESX-1 regulation and host membrane damage

Beyond its structural consequences, perturbation of envelope homeostasis is expected to impact regulatory networks that coordinate virulence-associated secretion and host interaction. In mycobacteria, envelope composition, lipid metabolism, and ESX-1 activity are tightly interconnected through regulatory circuits involving PhoPR, EspR, and WhiB6, which coordinate expression of secretion substrates with cell-wall and lipid biosynthetic pathways (Bosserman Rachel and Champion Patricia, 2017, Blasco et al., 2012, Koleske et al., 2025, Canestrari et al., 2025, Murdoch et al., 2026). Interestingly, Mm Δ*facl6* exhibited an altered ESX-1 transcriptional profile already under axenic conditions and did not show the coordinated ESX-1 induction observed in Mm wt during the intravacuolar phase of infection. In Mm wt, ESX-1-associated genes were strongly induced during residence within the MCV, consistent with a compartment-specific requirement for membrane damage to facilitate cytosolic access. By contrast, elevated basal ESX-1-associated transcription in Mm Δ*facl6* may contribute to the enhanced and premature membrane damage observed early during infection, while the absence of sustained ESX-1 induction at later stages suggests a dysregulation of the secretion system once bacteria exit the vacuole. Notably, dynamic regulation of ESX-1 in response to sustained stress has been reported in other contexts. For example, under nitric oxide exposure in Mm, WhiB6 was shown to initially activate ESX-1 expression before progressively repressing the locus while maintaining induction of stress-adaptive programs, including the DosR regulon (Chen et al., 2016). This demonstrates that mycobacteria can temporally modulate ESX-1 engagement, as observed upon cytosolic escape in our case.

At the cellular level, these transcriptional alterations correlated with enhanced and sustained host membrane damage responses. Cells infected with Mm Δ*facl6* showed increased recruitment of ESCRT components, Galectin-3 exposure, ubiquitination of bacteria, and LC3 association, indicating more frequent or prolonged disruption of the MCV membrane. Importantly, although Mm Δ*facl6* bacteria gained cytosolic access earlier and more efficiently than Mm wt, this did not favor their intracellular growth, underscoring that cytosolic exposure *per se*, contrary to Mm wt, is not sufficient to confer a fitness advantage to Mm Δ*facl6* in the absence of appropriate metabolic adaptation.

### Intra-host compartmentalization dictates lipid acquisition programs in intracellular mycobacteria

The compartment-specific transcriptional signatures observed in intracellular Mm indicate that lipid utilization is tightly constrained by the localization of the bacteria inside their host. Dual RNA-seq revealed that genes involved in sterol uptake and catabolism (i.e. Mce4 and KstR-regulated genes) were strongly induced when bacteria reside within the MCV, whereas this induction is attenuated upon cytosolic access. Sterol-associated transcriptional programs have been widely reported to be upregulated during macrophage infection by pathogenic mycobacteria and to contribute to intracellular survival and virulence, particularly in Mtb (Pandey and Sassetti, 2008, Griffin et al., 2012, Rienksma et al., 2018). Our findings extend these observations by demonstrating that sterol metabolic gene induction correlates with the intravacuolar phase of the infection. This compartment-specific regulation suggests that engagement of sterol utilization reflects exposure to the neutral lipid-rich intravacuolar environment (Barisch et al., 2015b, Perret et al., 2026) rather than infection status alone. Similarly, a subset of FAAL/FACL family genes (including *facl6*) were upregulated in intravacuolar Mm *wt* but showed little induction in the Mm *Δfacl6* mutant or in bacteria that prematurely escape to the cytosol. In line with this model, a recent preprint shows that loss of ESX-1 functions in Mtb reduces neutral lipid accumulation within intracellular bacteria during macrophage infection, implicating MCV damage as a prerequisite for access to host lipid pools (Padilla-Gomez et al., 2025). Together, these data indicate that intracellular compartmentalization dictates lipid availability over time and, in turn, shapes the metabolic programs engaged by intracellular mycobacteria.

### FACL6 enables metabolic buffering of host lipid fluctuations during intracellular infection

The compartment-dependent lipid programs observed in intracellular bacteria raise the question of how mycobacteria tolerate fluctuations in lipid availability and composition across infection stages and intracellular compartments. Such tolerance is thought to rely, at least in part, on fatty acid activation systems that couple lipid uptake to downstream metabolic fates, thereby preventing uncontrolled accumulation of potentially toxic lipid intermediates (Li et al., 2010, Weimar et al., 2002). In mycobacteria, fatty acyl-CoA synthetases (FACLs) convert free fatty acids into acyl-CoA thioesters that can be routed toward catabolism, neutral lipid storage, or cell-envelope biosynthesis, effectively buffering lipid flux under changing environmental conditions (Trivedi et al., 2004, Mohanty et al., 2011). Among them, FACL6 has been implicated in long-chain fatty acid activation and vectorial acylation, and its disruption In Mtb reduces TAG accumulation under stress conditions (Daniel et al., 2014, Mater et al., 2022). Our *in vitro* analyses extend these findings by showing that absence of FACL6 does not abolish fatty acid utilization, but instead alters how mycobacteria respond to changes in lipid abundance. The Mm Δ*facl6* mutant tolerated nutrient-poor conditions and moderate fatty acid concentrations, yet became acutely sensitive to high levels of saturated fatty acids. These observations indicate that FACL6 influences the lipid concentrations at which fatty acids transition from a growth-supporting to a growth-inhibitory input, consistent with a role in buffering excess lipid flux rather than directly controlling uptake.

The relevance of such buffering becomes particularly evident in intracellular contexts, where lipid exposure is imposed by host cell physiology and intracellular compartmentalization. Host LDs have been shown to accumulate at the MCV and serve as a source of neutral lipids that are transferred into intracellular bacteria and stored as TG within ILI (Barisch et al., 2015a, Barisch and Soldati, 2017). Manipulating host lipid metabolism in this system strongly alters bacterial lipid storage. Beyond serving as static storage depots, ILI are increasingly viewed as dynamic organelles that participate in regulating lipid flux, balancing TG synthesis and hydrolysis in response to metabolic state (Mallick et al., 2021, Maurya et al., 2019). In this framework, efficient activation of fatty acids would be required not only for *de novo* TG synthesis but also for reutilization of fatty acids released during ILI turnover (Fig. 1A). This raises the possibility that FACL6 contributes to buffering intracellular lipid flux by supporting fatty acid activation during periods of high TG turnover. Consistent with the importance of this host-pathogen lipid interface, recent work in macrophage models has shown that impairing host fatty acid import or storage restricts Mtb replication, highlighting how host lipid handling directly shapes pathogen fitness (Simwela et al., 2025).

### Intracellular growth depends on compartment-specific lipid processing

If lipid exposure differs across compartments, defects in lipid handling should become most apparent under conditions of high lipid flux. In this context, the behavior of the Mm Δ*facl6* mutant is particularly informative. Although Mm Δ*facl6* gains premature access to the cytosol, its intracellular growth is not restored to Mm wt levels, even in Dd *atg1*-KO cells where bacteria undergo an extended cytosolic phase. This dissociation indicates that, beyond enhanced membrane damage and premature cytosolic exposure, additional constraints limit the intracellular fitness of Mm Δ*facl6*. One plausible contributor is metabolic stress linked to lipid consumption. Consistent with this idea, our dual RNA-seq revealed that infection with Mm Δ*facl6* elicits a distinct host transcriptional response enriched for lipid metabolic pathways, including fatty acid metabolism, sterol-associated processes, and CoA biosynthesis, consistent with altered host lipid handling during the early divergence phase, which could reshape lipid pools available for mycobacteria. In this context, the intrinsic fatty acid sensitivity of Mm Δ*facl6* observed *in vitro* becomes particularly relevant, as the mutant displays a narrowed tolerance window for saturated fatty acid exposure.

Experimentally increasing host lipid load supported this compartment-centric view. Supplementation of host cells with excess palmitate, which promotes host neutral lipid accumulation (Du et al., 2013), resulted in a pronounced intracellular growth defect for both Mm wt and Mm Δ*facl6* in Dd wt cells. In contrast, this inhibitory effect was strongly attenuated in the Dd *atg1*-KO background, where infection is biased toward an early and sustained cytosolic exposure. Notably, palmitate supplementation did not further restrict growth of Mm Δ*facl6* in *atg1*-KO cells. Together, these results indicate that lipid overload is most detrimental in infection contexts dominated by intravacuolar phases, and that the impact of excess host-derived lipids is mitigated when bacteria escape early to the cytosol. Combined with the *in vitro* fatty acid sensitivity phenotypes, these findings suggest that FACL6-dependent lipid handling becomes particularly relevant when bacteria experience high lipid flux within the MCV, where excess fatty acids may need to be buffered into compatible metabolic or storage states to sustain intracellular growth.

### When and where do mycobacteria process host-derived lipids?

Bringing these observations together, our data suggest that the benefit of host-derived lipids is conditional on when and where bacteria encounter them and on their capacity to route lipid influx into compatible metabolic and storage states. When residing within the MCV, bacteria engage robust sterol acquisition programs and display greater neutral lipid storage signatures, whereas cytosolic access correlates with attenuated induction of sterol-associated genes and reduced lipid storage. In Mm Δ*facl6*, premature escape from the MCV occurs alongside impaired engagement of compartment-linked lipid programs, and the *in vitro* lipid sensitivity phenotypes provide a mechanistic rationale for why increased cytosolic access is not sufficient to restore growth: adaptation requires not only access to nutrients, but also the capacity to tolerate and process lipid pools imposed by distinct intracellular compartments.

## Acknowledgments

We gratefully acknowledge Dr. Dimitri Moreau, Dr. Stefania Vossio and Dr. Vincent Mercier, from the ACCESS Geneva Imaging Facility (University of Geneva) for their assistance and advice with high-content microscopy imaging and segmentations. A special thanks to Jérôme Bosset, from the Photonic Bioimaging Center (University of Geneva) for his help in SEM images acquisition. We gratefully acknowledge the Integrated Bioimaging Facility (iBiOs) at the University of Osnabrück and the Advanced Light and Fluorescence Microscopy (ALFM) Facility (University of Hamburg) at the CSSB. We thank Carola Schneider and Dr. Rudolph Reimer at the Leibniz Institute of Virology (LIV) for their support with EM image acquisition, and Dr. Aby Anand for acquiring the images. We are grateful to Dr. Gareth Prosser (Research Center Borstel) and Prof. Dr. Katia Cosentino (University of Modena and Reggio Emilia) for valuable advice and support. We deeply acknowledge Anthony Orvedahl (Washington University School of Medicine) for generously sharing the BV-2 *atg5*-KO cell lines. We thank Dr. Nabil Hanna for his assistance in coordinating the sequencing of the samples. We acknowledge the receipt of reagents and bacteria species from BEI Resources, NIAID, NIH, including purified PGL and PDIM used in this work, and Prof. Dr. Petry Broz for sharing the iBMDMs. This work was supported by the Swiss National Science Foundation grants 310030_188813 and 310030_219364 (TS) as well as the SystemsX.ch project HostPathX (HH, TS). Funding was provided by the German Research Foundation (DFG) within the SPP2225 (CB and iBios EM facility) and the projects SFB1557-P1 (CB) as well as SFB1557-Z02 (iBios EM facility). An EMBO post-doctoral fellowship was awarded to MF (ALTF 715-2021). Work in DS-lab is supported by the German Centre of Infection Research (TTU-TB).

## Material and methods

### Dd cell culture

Dd wt or *atg1*-KO (AX2) were grown axenically at 22°C in HL5-C medium (Formedium) containing 100 U/mL penicillin and 100 µg/mL streptomycin (P/S). Plasmids were electroporated into Dd and selected with the appropriate antibiotic. Hygromycin was used at a concentration of 50 mg/ml and Blasticidin at a concentration of 5 mg/ml.

### Mammalian cell lines and culture

#### • L929 cell culture

Cells were thawed into a small flask, expanded into medium flasks, and maintained in culture for two weeks. Before each trypsinization, the supernatant was collected, filtered through a 0.22 µm membrane, and stored at 4 °C; additional flasks were set up to increase yield. When ∼500 ml of supernatant had been accumulated, all batches were pooled, aliquoted into 50 ml tubes, and frozen at −80 °C. The resulting M-CSF–conditioned medium is stable for 3–4 months at −80 °C.

#### • iBMDM and BV-2 cell culture

iBMDMs were recovered by retrieving cryovials from liquid nitrogen, keeping them briefly on ice, and thawing them in a 37 °C water bath until just melted. Cells were transferred to conical tubes, slowly diluted dropwise with pre-warmed plain Dulbecco’s Modified Eagle Medium (DMEM), centrifuged (5 min, 170 × g), and the pellet was resuspended in iBMDM medium supplemented with P/S. Cells were plated in non–tissue-culture–treated Petri dishes and incubated at 37 °C with 5% CO₂. For routine passaging, cultures were chilled at 4 °C for 10–15 min to promote detachment, collected by gentle scraping, pelleted (5 min, 170 × g), and resuspended in fresh iBMDM + P/S medium. Cells were re-plated at appropriate density, typically at a 1:20 split ratio (or 1:5–1:6 from confluent cultures), and excess cells were discarded. For experimental seeding, cells were harvested as described, counted using the Countess II (Thermofisher), and resuspended at the required concentration in iBMDM medium before plating. For cryopreservation, harvested and counted cells were pelleted, resuspended in heat-inactivated FCS, and adjusted to 1 × 10⁷ cells/mL in FCS containing 10 % DMSO. Aliquots (1 mL) were dispensed into labelled cryovials, frozen at -80 °C in an insulated container, and transferred to liquid nitrogen after 24–48 h. BV-2 cells were grown at 37°C in DMEM complemented with 10% fetal bovine serum (FBS) and P/S.

### Bacterial strains and growth conditions

*Escherichia coli* Top10, DH5α and HB101 were used for cloning and grown at 37°C in LB medium. Mm was cultured in 7H9 or 7H12 media supplemented with 10 % OADC or ADC, 0.05% tyloxapol at 32 °C with shaking at 150 rpm. For growth assays under defined carbon source conditions, Mm cultures were pre-grown for 24 h in 7H9 medium to an OD₆₀₀ of 0.8-1.0 at 32°C with shaking (150 rpm). Cultures were harvested by centrifugation (3000 rpm, 10 min), washed, and resuspended in 7H12 medium. Bacterial suspensions were mechanically dispersed by passage through a 25-gauge needle (10 strokes) to reduce clumping and adjusted to the desired volume. 7H12 medium was supplemented with cholesterol, palmitate, or oleate at the indicated final concentrations. A 500 mM cholesterol stock solution was prepared in methanol:tyloxapol (1:1, v/v) by dissolving cholesterol in methanol at 80°C with intermittent vortexing for 20 min, followed by addition of tyloxapol and incubation at 80°C for an additional 10 min. The stock solution was stored at 4°C. An intermediate 2 mM cholesterol solution was prepared in pre-warmed 7H12 medium to facilitate solubilization prior to addition to cultures. Palmitate and oleate stock solutions (100 mM) were prepared similarly in methanol:tyloxapol (1:1, v/v) and diluted directly into 7H12 medium to the desired final concentration. Defined media were inoculated at a final density of 2.5x10⁶ bacteria/mL. Cultures were incubated at 32°C with agitation (150 rpm) for 48-72 h.

### Generation of Δ*facl6* mutant and molecular modifications

#### • Mm Δ*facl6* mutant generation

Mm Δ*facl6* was generated by specialized phage transduction as previously described (Koliwer-Brandl et al., 2016) with modifications. The flanking regions of *facl6* were amplified using the primer pairs oHK210/oHK211 and oHK212/oHK213. The PCR products were digested with AlwNI and Van91I, respectively, and ligated with Van91I-digested p0004S vector fragments. The resulting plasmids pHK65 and the DNA of the temperature-sensitive phage phAE159 were linearized with PacI before ligation. High-titer phages were prepared in *M. smegmatis* to perform specialized transduction in Mm to yield Δ*facl6* (Hyg). Then, the hygromycin-resistant *facl6* mutant was unmarked by applying the phAE7.1 phage, which contains the γδ-tnpR gene that produces the resolvase. The mutants obtained were verified by PCR using primer pairs with one primer located in the genome (facL6-L/-R-ext) and the other primer located in the resistance cassette (oHK33 and oHK36) and sequencing of the amplicon.

#### • Electroporation conditions

A 10-mL culture of Mm grown in 7H9 medium to OD >> 1 was harvested and washed twice in freshly prepared 10% glycerol containing 0.05% Tyloxapol at room temperature (3000 rpm, 10 min). The pellet was resuspended in 1 mL of the same buffer, and 100-µL aliquots were prepared for electroporation. For each transformation, 1-5 µg of plasmid DNA (thoroughly desalted) was pipetted directly into a pre-chilled 0.2-cm gap Bio-Rad electroporation cuvette. Subsequently, 100 µL of washed bacterial suspension was added and gently mixed. Electroporation was performed using a Bio-Rad Gene Pulser with settings of 2.5 kV, 1 kΩ, and 25 µF. Immediately after the pulse, 250–1000 µL of 7H9 medium was added to the cuvette, and cells were transferred to a tube and allowed to recover at 30 °C for at least 4 h. Following recovery, 50–200 µL of the culture was plated on selective 7H10 agar.

#### • Cloning of genetic complementation and reporter plasmids

For genetic complementation, the flanking regions of *facl6* were amplified using the primer pairs oHK263/oHK260 or oHK259/oHK260 from genomic DNA to introduce the restriction sites HindIII and/or PacI. The purified PCR product was cloned into the plasmid backbone pMV361-PacI (Koliwer-Brandl et al., 2016), a plasmid that integrates in the attB site of mycobacterial genome, gaining the plasmid pHK84. For the generation of a FACL6 reporter plasmid, the natural promoter region of *facl6*, about 500 bps downstream of facl6, was cloned into the mCherry expressing plasmid mCherry. To this end, the promoter region was amplified using the primer pair oHK274/oHK244 and genomic DNA as template introducing the restriction cleavage sites PacI and XbaI. To insert a PacI cleavage site, the mCherry gene was amplified using the primers oHK272/oHK273 from the plasmid and the PCR product was digested with HindIII and PacI. The two PCR products were then cloned into XbaI/HindIII-digested pCherry8 gaining pHK87. The resulting strain was then transformed with pmsp12::GFP (Addgene #30171). eBFP, GFP, mCherry and luciferase expressing bacteria were generated by transforming the unlabelled Mm strains wt, Δ*facl6*, Δ*facl6:: facl6* and ΔRD1 with the pTEC18 plasmid (addgene #30177) the pMSP12::DsRed/GFP plasmid (addgene #30167), the pCherry10 plasmid (addgene #24664) and the pMV306hsp+LuxG13 plasmid expressing luciferase (addgene #26161) (Andreu et al., 2010), respectively, and grown in medium with 100 µg/ml hygromycin or 25 µg/ml kanamycin. Fluorescent transformants typically appeared after ∼10 days, showing characteristic coloration depending on the fluorophore (e.g., green for GFP, red for mCherry).

### Minimum Inhibitory Concentration (MIC) assay

Resistance against antimicrobial compounds was determined as described previously (Mulholland et al., 2024) using the microbroth dilution method and luciferase-expressing Mm strains. To this end, two-fold serial dilutions of the drugs were prepared in 7H9/OADC/glycerol/tyloxapol in white U-bottom 96-well plates. Luciferase-expressing Mm strains were inoculated two days before the assay.

Cultures were diluted the day before to reach an OD₆₀₀ of approximately 1.0 on the day of the experiment. The bacterial suspensions were diluted to obtain an OD₆₀₀ of 0.002 (equivalent to 1 × 10⁵ cells/ml) and transferred to each well resulting in a final well volume of 100 μl per well with compounds at the desired concentrations. Plates were sealed and incubated at 32°C for 4 days. Luminescence was recorded using a Tecan plate reader with an integration time of 2500 ms. For data analysis, relative growth was calculated by normalizing each well’s relative light units (RLU) to the mean of the no-drug control (100% growth) and the mean of the Rifampicin control (0% growth). Percentage growth was computed as:

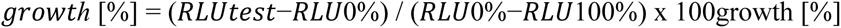

Dose–response curves were generated using GraphPad Prism. IC₅₀ values were obtained by nonlinear regression using the “[Inhibitor] vs. response – variable slope (four-parameter)” model, constraining the top plateau to 100 and the bottom plateau to values greater than –1.

### Promotor activity assay by flow cytometry

FACL6 promotor activity *in vitro* was assessed by flow cytometry. Briefly, bacteria were washed twice in PBS and syringed. 10,000 cells and 20,000 bacterial events per condition were analysed using a SonySH800. Flow cytometry plots were generated with Cell Sorter Software by Sony. mCherry fluorescence served as the promoter-dependent signal, while bacteria were identified by constitutive GFP expression. For all analyses, the ratio of mCherry to GFP fluorescence (mCherry/GFP) was used to quantify promoter activity.

### Lipid extraction and TLC

Mm lipids were extracted with chloroform:methanol (1:2, v/v) as described previously (Bligh and Dyer, 1959). After lipid extraction, lipid samples were re-suspended in 100µL chloroform (for a protein content in the initial sample equivalent to 1 mg/mL) and spotted on silica high performance TLC plates. Glycolipids were separated in a system containing chloroform:methanol (60:40, v:v), and PDIM and neutral lipids were separated with a system consisting of petroleum ether:diethylether (90:10, v:v). Both systems ran to 1cm below the upper border. After drying plates under a fume hood and total lipids were detected by charring with a solution containing 0.63g MnCl2.4H2O, 60mL water, 60mL methanol and 4mL concentrated sulfuric acid). Lipid standards were kindly obtained from BEI Resources: purified PGL from *M. canettii* (NR-36510) and purified PDIM from *M. tuberculosis* (NR-20328).

### Dd infection assay

Dd were grown in HL5c-filtered medium (without penicillin/streptomycin) to > 80% confluency on 10-cm dishes (a confluent dish corresponding to ∼5 × 10⁷ cells). For infection, cells were overlaid with 3.5 mL HL5c-filtered medium. Mm strains were cultured for at least 24 h in 7H9/OADC at 32 °C to OD₆₀₀ 0.8 – 1.0. For MOI 10 per 10 cm dish, 5 × 10⁸ bacteria were pelleted (12,000 rpm, 4 min), washed once in HL5c-filtered medium, and resuspended in 500 µL. Bacteria were declumped by passage through a blunt 25-gauge needle (∼10 strokes) under microscopic control. The bacterial suspension was added directly onto the amoeba monolayer, and plates were sealed with parafilm and centrifuged twice at 500 × g for 10 min at room temperature, gently redistributing the inoculum between spins. Cells were allowed to phagocytose bacteria for an additional 20 min before removal of extracellular bacteria by 3–8 washes with HL5c filtered medium. Infected cells were detached by tapping, collected in HL5c filtered medium supplemented with low-dose P/S (1:2000), and resuspended to the required volume depending on the downstream experiment. The intracellular growth was monitored by measuring the luminescence signal with a Synergy Mx Monochromator-Based Multi-Mode Microplate Reader (Biotek) at 25°C in a white 96-well plate.

### iBMDM infection assay

One day prior to infection, 1–5 × 10⁶ iBMDMs were seeded per well in 6-well plates (2 mL medium/well) and incubated at 37 °C, 5% CO₂. Additional wells were plated for cell counting at the day of infection. For bacterial preparation, mycobacterial strains were pre-cultured for 48 h in 7H9 medium supplemented with OADC and appropriate antibiotics. For the infection, Mm cultures were grown to an OD₆₀₀ ≈ 0.8–1.0. Macrophage confluence and viability were confirmed microscopically, and viable cell numbers were determined by trypan blue staining. Cell number was determined with an automated cell counter (Thermofisher). Bacterial cultures were measured at OD₆₀₀ and diluted to achieve a MOI of 5. Bacteria were pelleted (12,000 rpm, 4 min), washed once in DMEM, resuspended in iBMDM medium, and mechanically dispersed by repeated passage through a 25-gauge needle. After declumping, bacteria were added to the cells, plates were sealed with parafilm, and centrifuged twice at 500 × g for 15 min at room temperature to synchronize uptake. Cells were allowed to phagocytose for 30 min at 37 °C, washed four times with PBS, detached using 2 mM EDTA, and resuspended in DMEM containing 10% FBS and 0.5% penicillin/streptomycin. For intracellular growth assay, infected iBMDM were stored at 32°C and luminescence measured around 3-4 times per day. Luminescence was measured by Infinite 200 pro M-plex plate reader (Tecan).

### BV-2 infection assay

The same protocol as for Dd infection was followed for BV-2 infections, with slight modifications. The day before the infection, 5 × 10^5^ cells/ml were plated in DMEM complemented with FBS. Cells were infected at a MOI of 5 and maintained at 32°C. Infection was monitored by microscopy with a 40× objective with the ImageXpress Micro XL HC microscope every 1.5 hours for 36 hours using three wells per condition with nine imaged fields per well with three sections per field with a 1-μm interval. Analysis was performed with the MetaExpress software and quantifying area of the GFP signal.

### RT-qPCR

Total RNA of cells infected with Mm wt was extracted at the indicated time points using the Direct-zol RNA MiniPrep kit (Zymo Research) following the manufacturer’s instructions. RNA (1 μg) was retrotranscribed using the iScript cDNA Synthesis Kit and polydT primers (Bio-Rad). The cDNA was amplified and the SsoAdvanced universal SYBR Green supermix (Bio-Rad). Amplimers for *facl6* and *gapdh* were detected on a CFX Connect Real-Time PCR Detection System (Bio-Rad). The housekeeping gene *gapdh* was used for normalization. PCR amplification was followed by a DNA melting curve analysis to confirm the presence of a single amplicon. Relative mRNA levels (2^−ΔΔCt^) were determined by comparing first the PCR cycle thresholds (Ct) for *facl6* and *gapdh* (ΔC) and, second, Ct values in infected cells versus mock-infected cells (ΔΔC).

### RNA-sequencing

Following infection with GFP-expressing bacteria, the cell population was pelleted and resuspended in 500 μl of HL5c and sorted by FACS (Beckman Coulter MoFlo Astrios). The gating was based on cell diameter (forward scatter) and granularity (side scatter). The infected (GFP-positive) sub-fraction was selected based on the GFP intensity (FITC channel). Mock-infected cells were subjected to the same treatments and collected as GFP negative. Typically, ∼ 5 × 10^5^ cells of each fraction were collected for RNA isolation. RNA isolation was performed as for qRT-PCR. Quality of RNA libraries, sequencing, and bioinformatic analysis were performed as previously described (Hanna et al., 2019, Raykov et al., 2023, Perret et al., 2026).

### Live imaging

To monitor non-infected cells or the course of infection by spinning disc microscopy, cells were transferred to either 4- or 8-well m-ibidi slides and imaged in low fluorescent medium (LoFlo, Formedium, UK) with a Zeiss or Nikon Cell observer SD microscope using the 63x oil objective (NA 1.46). Z-stacks of 15-20 slices with 300 nm intervals were acquired. The images were further processed and analysed with ImageJ. To analyse plasma membrane damage, wells with cells were washed with PBS and covered with transparent DMEM containing 0.5 µM pyromarker ToPro3TM-iodide (Invitrogen).

### Super-Resolution via Optical Re-Assignment (SoRa) of Mm

Mm strains were inoculated in fresh 7H9 medium supplemented with ADC or OADC and grown to an optical density of OD₆₀₀ = 1.0. Cultures were then diluted to OD₆₀₀ = 0.5 in fresh medium and incubated for 24 h. After washing with PBS. pellets were resuspended in 3 % paraformaldehyde (PFA) and fixed for 30 min at room temperature on a shaker, followed by a gentle PBS wash. Fixed cells were stained with 10 µM Bodipy 493/503 for 60 min at room temperature and washed twice with PBS and once with ddH₂O. To disperse aggregates, bacteria were gently syringed 8–10 times through a 25-gauge blunt needle. After embedding (ProLong, Invitrogen), the mixture was placed on a microscope slide, and a poly-L-lysine–coated coverslip was gently placed on top to evenly spread the bacteria. Mounted samples were protected from light to prevent photobleaching and imaged immediately using a Nikon spinning disc microscope and SoRa.

### Antibodies, fluorescent probes and immunofluorescence

The anti-LC3 antibody (BLD-125410) and the AF488-conjugated anti-mouse/human Mac-2 (Galectin-3) antibody (BLD-125410) were obtained from Biozol (BioLegend). The Adipophilin (PLIN2) antibody (FGI-20R-AP002-100UL) from Biozol (Fitzgerald). The anti-Ub FK2 antibody (ENZ-ABS840) was obtained from Enzo Life Sciences. Goat anti-rabbit, goat anti-mouse, and goat anti-rat IgG secondary antibodies conjugated to Alexa Fluor 546 (Thermo Fisher Scientific) or to the CF488R, CF568, and CF640R fluorophores (Biotium) were used. Bodipy493/503 was used to stain neutral lipids of Mm.

For immunostaining, cells were seeded on acid-cleaned poly-L-Lysine coated 10 mm coverslips and centrifuged at 500 g for 10 min at RT. Cells were fixed with 4 % PFA/ picric acid (Dd) or 4% PFA (iBMDMs). For Dd, coverslips were washed twice for 15 min in PBG (PBS + 0.2% gelatin) and incubated with primary antibodies diluted in PBG overnight at 4 °C. After six 5-min washes in PBG, secondary antibodies diluted in PBG were applied for 1.5 h at room temperature, followed by two 5-min washes in PBG and three 5-min washes in PBS. Coverslips were briefly dipped in ddH₂O, excess fluid removed, and mounted in antifade medium on glass slides, sealed with nail polish, and dried for 24 h at room temperature before storage at 4 °C. For antibody staining of iBMDM, fixed cells on coverslips were washed with PBS and incubated for 30 min in permeabilization buffer (0.05g saponin and 0,1g BSA dissolved in 50mL PBS). Primary antibodies were diluted to the required concentration in permeabilization buffer. Coverslips were briefly dried, placed cell-side up on a parafilm-coated moist chamber, and incubated with 50 µL of the diluted primary antibody for 60–90 min at room temperature. After three washes with permeabilization buffer, cells were incubated with the appropriate secondary antibodies (diluted in permeabilization buffer) for 1 h at room temperature in the dark; diluted secondary antibodies were centrifuged at maximum speed for 5 min at 4 °C prior to use to remove aggregates. Coverslips were subsequently washed three times with permeabilization buffer and twice with PBS. For embedding, a drop of embedding medium was placed on a microscope slide, excess liquid was removed from coverslips, and coverslips were mounted cell-side down onto the medium. Samples were dried O/N at 4 °C. Images were acquired using an Olympus LSM FV3000 NLO microscope with a 60x oil objective (NA of 1.40) and Nikon SD confocal microscope 63x oil objective, respectively. 10-20 slices with 300 nm intervals were taken. All images were analysed and processed using ImageJ and image quantifications were generated using Graphpad Prism.

### Electron microscopy (EM)

A confluent 6-cm culture dish of infected Dd cells was fixed for 1 h in HL5C medium containing 2% glutaraldehyde. Samples were then stained for 20 min with 2% osmium tetroxide buffered in 0.1 M imidazole as previously described (Angermuller and Fahimi, 1982). Cells were detached using a scraper, transferred into 1 ml PBS in a microcentrifuge tube, and washed twice with PBS. The samples were subsequently submitted to the EM facility at the University of Geneva, Faculty of Medicine for further preparation. There, samples underwent postfixation, embedding in Epon resin, and standard ultrathin section processing as previously described (Orci et al., 1973). Imaging was performed using a Tecnai 20 transmission electron microscope (FEI).

### Serial block-face (SBF) - scanning electron microscopy (SEM)

For SBF-SEM analysis, Dd cells infected with Mm were harvested 24 hours post-infection. Fluid-cultured, infected cells were transferred to falcon tubes and fixed by adding 3 ml of 4% glutaraldehyde in HL5C medium to 3 ml of cells in HL5C medium (final concentration 2% glutaraldehyde) for 30 minutes at room temperature. Cells were centrifuged at 250G for 3-5 minutes to form pellets, washed in HL5C medium, transferred to fresh Eppendorf tubes, and washed in 0.1 M imidazole buffer. Sample processing was performed according to a modified NCMIR rOTO-post-fixation protocol (Deerinck et al., 2010), which provides enhanced contrast and resistance to electron beam damage during serial imaging of specimens embedded in hard EPON resin. In brief, post-fixation was performed in 4% osmium tetroxide (Science Services) combined with 3% potassium ferrocyanide in 0.2 M imidazole buffer, diluted in 0.1 M imidazole buffer to achieve final concentrations of 2% osmium tetroxide and 1.5% potassium ferrocyanide, for 1 hour on ice. All incubation steps were performed with gentle agitation. Unless otherwise specified, samples were washed five times (3 minutes each) in ddH₂O at room temperature between each step. Cells were subsequently exposed to 1% (w/v) thiocarbohydrazide (Sigma Aldrich) preheated to 60°C in ddH₂O for 20 minutes at room temperature. A second osmication step was performed using 2% osmium tetroxide in water for 30 minutes at room temperature. Additional contrast enhancement was achieved through incubation in 1% tannic acid (Sigma Aldrich) in ddH₂O for 30 minutes at room temperature. Samples were stained with 1% aqueous uranyl acetate (EMS) overnight at 4°C, then warmed to approximately 50°C prior to washing. Five washes with pre-heated water (∼50°C, 3 minutes each) were performed. The final en bloc staining step consisted of incubation in freshly prepared Walton’s lead aspartate solution (Lead(II) nitrate (Carl-Roth), L-Aspartate (Serva) at pH 5.5) for 30 minutes at 60°C. Dehydration was performed using a graded ethanol series (50%, 70%, 90%, and two rinses in 100% ethanol) on ice, 5 minutes per step, followed by two rinses in anhydrous acetone (VWR Chemicals), the first on ice and the second at room temperature (10 minutes each). Resin infiltration was carried out using increasing ratios of Epon:acetone (3:1, 1:1, 1:3) for 2 hours per step, with a subsequent overnight incubation in pure Epon 812 (Sigma Aldrich). To ensure proper sample conductivity, the final infiltration steps were performed using resin containing 1.5% (w/w) carbon filler Ketjenblack (TAAB). Polymerization occurred in hard Epon within Eppendorf tubes at 60°C for 48 hours. After polymerization, the pellet-containing bottom portion of each Eppendorf tube was excised from the resin block. Samples were trimmed to dimensions appropriate for SBF-SEM imaging and mounted onto aluminum rivets using two-component conductive silver epoxy (Microtonano). A 20 nm gold coating was applied to enhance conductivity. The rivet-mounted sample was loaded into a 3View2XP Gatan stage (Gatan, Pleasanton, CA, USA) installed in a Jeol JSM 7200F scanning electron microscope (JEOL Ltd., Tokyo, Japan) and positioned parallel to the diamond knife edge. Optimal imaging conditions were established at 1.5 kV accelerating voltage, high vacuum mode (10 Pa), with a 30 nm condenser aperture and +550 V stage bias. Data acquisition parameters included pixel size 5 nm, 3.5 μs dwell time, slice thickness of 50 nm, and image dimensions of 12000×12000 pixels. Imaging continued until approximately 60×60×25 μm volumes were captured or until adequate numbers of complete cells were recorded. Data collection was managed using Gatan Digital Micrograph software (Version 3.32.2403.0). Post-acquisition processing, including stack alignment, filtering, and manual segmentation, was conducted in Microscopy Image Browser (MIB, Version 2.84) (Belevich et al., 2016). To this end, individual infected cells were cropped as regions of interest prior to analysis. Mycobacteria and infection-related structures were segmented in 3D using a combination of manual and semi-automated tools within MIB. Segmentation masks were refined by interpolation and morphological filtering, and all objects were verified in both 2D and 3D views. Quantitative measurements were extracted using MIB’s built-in voxel-based statistics tools. Final image stacks were binned to optimize computational efficiency before importing into Amira 3D (Version 2022.1, Thermo Fisher) for three-dimensional visualization and analysis.

### Software and Statistical Analysis

All microscopy images were analyzed and processed using FIJI/ImageJ or Imaris 9.5. (Bitplane, Switzerland). Co-localisation was quantified using either CellPathFinder (Yokogawa) or NIS-Elements AR software. In addition, a custom MATLAB program was developed for quantitative analysis of ILIs in live-cell imaging experiments. Graphs and statistical analyses were generated using GraphPad Prism 10. Data are presented as mean ± standard deviation (SD) or mean ± standard error of the mean (SEM), as indicated in the figure legends. Statistical significance was assessed using two-tailed, unpaired parametric t tests or two-way ANOVA followed by Fisher’s LSD test. Statistical significance is indicated as ns (not significant), P < 0.05 (*), P < 0.01 (**), P < 0.001 (***), and P < 0.0001 (****). “N” indicates the number of independent biological replicates, “n” indicates the number of technical replicates. AlphaFold-predicted structures were visualized and processed using PyMOL. Schemes and figures were generated with Biorender and Adobe Illustrator (2025 and 2026).

## Supplementary material

**Figure S1. Mm** Δ***facl6* has an intracellular growth defect and forms microcolonies in BV-2 and iBMDMs.** (**A**) Relative *facl6* expression (normalized to *gapdh*) in Mm wt, Δ*facl6*, and *Δfacl6::facl6* measured by RT-qPCR. (**B**) Phagocytic uptake of Mm strains by Dd, quantified at 3 hpi. (**C**) Intracellular growth of Mm wt, Δ*facl6*, Δ*facl6::facl6*, and ΔRD1 expressing GFP in BV-2 monitored by microplate reader over 48 h. (**D**) High-content images of infected BV-2 at 24 hpi (scale bars, 5 µm) (**E**) Intracellular growth of the same Mm strains in iBMDMs monitored by a microplate reader over 30 h. (**F**) Spinning disc images of infected iBMDMs at 48 hpi (scale bars, 5 µm). Growth curves show mean fold change ± SD (N=3) analyzed using a two-way ANOVA with Fisher’s LSD test (*p **<** 0.05, **p **<** 0.01, ****p **<** 0.0001, ns: not significant). RFU: relative fluorescence units. RLU: relative luminescence units.

**Figure S2. Mm** Δ***facl6* intracellular growth and microcolony formation in BV-2 is partially restored in absence of autophagy.** (**A)** BV-2 wt or *atg5*-KO cells were infected with Mm wt and Δ*facl6* expressing GFP and monitored for 36 hours using a microplate reader. Growth curves show mean fold change ± SD (N=3) analyzed using a two-way ANOVA with Fisher’s LSD test (****p **<** 0.0001). RFU: Relative Fluorescent Unit. (**B**) High-content images of infected BV-2 WT and *atg5*-KO at 24 hpi (scale bars, 5 µm). (**C**) The profile of DEGs related to the Esx-1 operon (previously defined gene lists) in Mm WT or Δ*facl6* grown in broth (7H9/OADC/glycerol/tyloxapol) was plotted as heatmap (each column represents an individual sample). Blue: downregulated genes, red: upregulated genes. (**D**) Esx-1 related genes differentially expressed between Mm wt or Mm Δ*facl6* grown in axenic medium. Blue: upregulated in Mm Δ*facl6*, red: downregulated in Mm Δ*facl6* compared to Mm wt.

**Figure S3. Complete dataset supporting Mm Δ*facl6*-induced MCV damage and cytosolic escape.** (**A-B**) Spinning disc images of Alix-GFP (A) or GFP-Vps32 (B) recruitment to MCVs in Dd infected with Mm wt, Δ*facl6*, Δ*facl6::facl6*, or ΔRD1, performed as described in Fig. 4. Samples were taken at the indicated time points. Arrows indicate Alix-GFP–positive or GFP-Vps32 puncta co-localizing with intracellular bacteria (scale bars, 5 µm). (**C**-**D**) Correlation between the number of Alix-GFP- and GFP-Vps32-positive puncta per cell and intracellular bacterial microcolony size. Data from all strains were pooled, and all microcolony size classes (including > 40 AU) are shown. Statistical significance was determined using an unpaired two-tailed Student’s t-test (*** p < 0.001, ns: not significant).

**Figure S4. Mm** Δ***facl6* induces exacerbated damage to its MCV in iBMDMs and is escaping earlier to the cytosol.** (**A**) Representative images of Galectin -3 (Gal3) recruitment to intracellular Mm in iBMDMs. (**B**) Quantification of Gal3-positive bacteria. (**C**) Representative images of Ubiquitin (Ub) recruitment to intracellular Mm. (**D**) Quantification of Ub-positive bacteria. (**E**) Representative images of PLIN2 recruitment to intracellular Mm. (**F**) Quantification of PLIN2-positive bacteria. (**G**) Representative images of LC3 recruitment to intracellular Mm. Arrows point to Gal-3, Ub-, PLIN2 or LC3-positive bacteria. iBMDMs were infected with Mm wt, Δ*facl6*, *Δfacl6::facl6*, or ΔRD1, fixed at the indicated time points and stained for the indicated host proteins. Images were acquired using a spinning disc microscope (scale bars, 5 µm). Quantifications were performed manually. Data represent mean ± SEM from three independent experiments, except for LC3 where two independent experiments were pooled. Statistical significance was determined using a two-way ANOVA with Fisher’s LSD test (*p **<** 0.05, **p **<** 0.01, ns: not significant).

**Fig. S5. Loss of FACL6 increases plasma membrane damage and host cell death in iBMDMs. (A)** Representative images of PyroMarker staining in iBMDMs infected with Mm wt, Δ*facl6*, Δ*facl6::facl6*, or ΔRD1 at 27 hpi. PyroMarker labels cells with compromised plasma membrane integrity. (**B**) Quantification of PyroMarker-positive, infected cells. (C) Representative images showing morphological features consistent with pyroptotic and necrotic cell death in infected iBMDMs at 3 hpi. (**D**) Quantification of pyroptotic and necrotic infected cells. iBMDMs were infected with the indicated Mm strains and stained at the indicated time points. Images were acquired using a spinning disc microscope (scale bars, 5 µm). Quantifications were performed manually. Data represent mean ± SEM from three independent experiments. Statistical significance was determined using a two-way ANOVA with Fisher’s LSD test (**p **<** 0.01, ns: not significant).

**Fig. S6. Transcriptomic changes in Mm *Δfacl6* inoculum and complete TLCs. (A)** Volcano plot showing differential gene expression between Mm Δfacl6 and wt grown in broth (7H9/OADC/glycerol/tyloxapol). Each dot represents one gene, plotted by log₂ fold change versus - log₁₀ *P* value. Significantly upregulated genes are shown in red, downregulated genes in blue, and non-significant genes in gray. Selected genes of interest are indicated. (**B**) Functional categorization of the top differentially expressed genes (DEGs). Pie charts show the distribution of upregulated and downregulated genes across functional categories. (**C-D**) Complete TLCs analyses corresponding to TLCs shown in Fig. 5D.

**Fig. S7 Complementing growth curves showing how FACL6 deletion alters Mm lipid sensitivity to host-lipids.** (**A**–**C**) Growth of GFP-expressing Mm wt in 7H12 medium supplemented with cholesterol (**A**), palmitate (**B**), or oleate (**C**). (**D**–**F**) Growth of GFP-expressing Mm Δ*facl6* under the same conditions. (**G-I**) Growth of GFP-expressing Mm Δ*facl6::facl6* under the same conditions. Experiments were performed as described in Fig. 6. Grey areas indicate the time window used for AUC calculations shown in Fig. 6. One representative experiment of three biological replicates is shown. Data represent mean ± SEM (n=3).

## References

Abdallah, A. M., Weerdenburg, E. M., Guan, Q., Ummels, R., Borggreve, S., Adroub, S. A., Malas, T. B., Naeem, R., Zhang, H., Otto, T. D., Bitter, W. & Pain, A. 2019. Integrated transcriptomic and proteomic analysis of pathogenic mycobacteria and their esx-1 mutants reveal secretion-dependent regulation of ESX-1 substrates and WhiB6 as a transcriptional regulator. PLoS One, 14, e0211003.

Alibaud, L., Rombouts, Y., Trivelli, X., Burguiere, A., Cirillo, S. L., Cirillo, J. D., Dubremetz, J. F., Guerardel, Y., Lutfalla, G. & Kremer, L. 2011. A Mycobacterium marinum TesA mutant defective for major cell wall-associated lipids is highly attenuated in Dictyostelium discoideum and zebrafish embryos. Mol Microbiol, 80, 919–34.

Anand, A., Mazur, A. C., Rosell-Arevalo, P., Franzkoch, R., Breitsprecher, L., Listian, S. A., Hüttel, S. V., Müller, D., Schäfer, D. G., Vormittag, S., Hilbi, H., Maniak, M., Gutierrez, M. G. & Barisch, C. 2023. ER-dependent membrane repair of mycobacteria-induced vacuole damage. *mBio*, e0094323.

Andreu, N., Zelmer, A., Fletcher, T., Elkington, P. T., Ward, T. H., Ripoll, J., Parish, T., Bancroft, G. J., Schaible, U., Robertson, B. D. & Wiles, S. 2010. Optimisation of bioluminescent reporters for use with mycobacteria. PLoS One, 5, e10777.

Angermuller, S. & Fahimi, H. D. 1982. Imidazole-buffered osmium tetroxide: an excellent stain for visualization of lipids in transmission electron microscopy. Histochem J, 14, 823–35.

Ates, L. S. 2020. New insights into the mycobacterial PE and PPE proteins provide a framework for future research. Mol Microbiol, 113, 4–21.

Aylan, B., Bernard, E. M., Pellegrino, E., Botella, L., Fearns, A., Athanasiadi, N., Bussi, C., Santucci, P. & Gutierrez, M. G. 2023. ATG7 and ATG14 restrict cytosolic and phagosomal Mycobacterium tuberculosis replication in human macrophages. Nat Microbiol, 8, 803–818.

Barisch, C., Lopez-Jimenez, A. T. & Soldati, T. 2015a. Live Imaging of Mycobacterium marinum Infection in Dictyostelium discoideum. Methods Mol Biol, 1285, 369–85.

Barisch, C., Paschke, P., Hagedorn, M., Maniak, M. & Soldati, T. 2015b. Lipid droplet dynamics at early stages of Mycobacterium marinum infection in Dictyostelium. Cell Microbiol, 17, 1332–49.

Barisch, C. & Soldati, T. 2017. Mycobacterium marinum Degrades Both Triacylglycerols and Phospholipids from Its Dictyostelium Host to Synthesise Its Own Triacylglycerols and Generate Lipid Inclusions. PLoS Pathog, 13, e1006095.

Beckwith, K. S., Beckwith, M. S., Ullmann, S., Sætra, R. S., Kim, H., Marstad, A., Åsberg, S. E., Strand, T. A., Haug, M., Niederweis, M., Stenmark, H. A. & Flo, T. H. 2020. Plasma membrane damage causes NLRP3 activation and pyroptosis during Mycobacterium tuberculosis infection. Nat Commun, 11, 2270.

Belevich, I., Joensuu, M., Kumar, D., Vihinen, H. & Jokitalo, E. 2016. Microscopy Image Browser: A Platform for Segmentation and Analysis of Multidimensional Datasets. PLOS Biology, 14, e1002340.

Bernard, E. M., Fearns, A., Bussi, C., Santucci, P., Peddie, C. J., Lai, R. J., Collinson, L. M. & Gutierrez, M. G. 2020. M. tuberculosis infection of human iPSC-derived macrophages reveals complex membrane dynamics during xenophagy evasion. J Cell Sci, 134.

Blasco, B., Chen, J. M., Hartkoorn, R., Sala, C., Uplekar, S., Rougemont, J., Pojer, F. & Cole, S. T. 2012. Virulence Regulator EspR of Mycobacterium tuberculosis Is a Nucleoid-Associated Protein. PLOS Pathogens, 8, e1002621.

Bligh, E. G. & Dyer, W. J. 1959. A rapid method of total lipid extraction and purification. Can J Biochem Physiol, 37, 911–7.

Block, A. M., Ravindran Nair, R., Meikle, V., Wiegert, P. C., White, D. W., Zhang, L., Niederweis, M. & Tischler, A. D. 2026. The Mycobacterium tuberculosis ESX-5 secretion system enables carbon source utilization and growth in mice. *mBio*, e0350025.

Boradia, V., Chen, J., Frando, A., Clark, L. V. & Grundner, C. 2025. PE/PPE proteins contribute to <EM>Mycobacterium tuberculosis</EM> drug resistance. bioRxiv, 2025.09.24.678375.

Bosserman Rachel, E. & Champion Patricia, A. 2017. Esx Systems and the Mycobacterial Cell Envelope: What’s the Connection? Journal of Bacteriology, 199, 10.1128/jb.00131-17.

Brennan, M. J. 2017. The Enigmatic PE/PPE Multigene Family of Mycobacteria and Tuberculosis Vaccination. Infect Immun, 85.

Broz, P., Von Moltke, J., Jones, J. W., Vance, R. E. & Monack, D. M. 2010. Differential requirement for Caspase-1 autoproteolysis in pathogen-induced cell death and cytokine processing. Cell Host Microbe, 8, 471–83.

Canestrari, J. G., Gordon, E. C., Bruce, S. A., Lasek-Nesselquist, E., Schultz, S. R., Marietta, H. A., Biegas, K. J., Swarts, B. M., Champion, M. M., Derbyshire, K. M. & Gray, T. A. 2025. WhiB6 Transduces Contact-Dependent Signaling in Mycobacterium smegmatis and Coordinately Induces Both ESX-1 and ESX-4. Mol Microbiol, 124, 559–572.

Cardenal-Munoz, E., Arafah, S., Lopez-Jimenez, A. T., Kicka, S., Falaise, A., Bach, F., Schaad, O., King, J. S., Hagedorn, M. & Soldati, T. 2017a. Mycobacterium marinum antagonistically induces an autophagic response while repressing the autophagic flux in a TORC1- and ESX-1-dependent manner. PLoS Pathog, 13, e1006344.

Cardenal-Munoz, E., Barisch, C., Lefrancois, L. H., Lopez-Jimenez, A. T. & Soldati, T. 2017b. When Dicty Met Myco, a (Not So) Romantic Story about One Amoeba and Its Intracellular Pathogen. Front Cell Infect Microbiol, 7, 529.

Chavadi, S. S., Edupuganti, U. R., Vergnolle, O., Fatima, I., Singh, S. M., Soll, C. E. & Quadri, L. E. 2011. Inactivation of tesA reduces cell wall lipid production and increases drug susceptibility in mycobacteria. J Biol Chem, 286, 24616–25.

Chen, Z., Hu, Y., Cumming, B. M., Lu, P., Feng, L., Deng, J., Steyn, A. J. & Chen, S. 2016. Mycobacterial WhiB6 Differentially Regulates ESX-1 and the Dos Regulon to Modulate Granuloma Formation and Virulence in Zebrafish. Cell Rep, 16, 2512–24.

Daniel, J., Sirakova, T. & Kolattukudy, P. 2014. An acyl-CoA synthetase in Mycobacterium tuberculosis involved in triacylglycerol accumulation during dormancy. PLoS One, 9, e114877.

Dargham, T., Mallick, I., Kremer, L., Santucci, P. & Canaan, S. 2023. Intrabacterial lipid inclusion-associated proteins: a core machinery conserved from saprophyte Actinobacteria to the human pathogen Mycobacterium tuberculosis. FEBS Open Bio, 13, 2306–2323.

DE Nardo, D., Kalvakolanu, D. V. & Latz, E. 2018. Immortalization of Murine Bone Marrow-Derived Macrophages. Methods Mol Biol, 1784, 35–49.

Deerinck, T., Bushong, E., Lev-Ram, V., Shu, X., Tsien, R. & Ellisman, M. 2010. Enhancing Serial Block-Face Scanning Electron Microscopy to Enable High Resolution 3-D Nanohistology of Cells and Tissues. Microscopy and Microanalysis, 16, 1138–1139.

Deretic, V., Saitoh, T. & Akira, S. 2013. Autophagy in infection, inflammation and immunity. Nat Rev Immunol, 13, 722–37.

Du, X., Barisch, C., Paschke, P., Herrfurth, C., Bertinetti, O., Pawolleck, N., Otto, H., Ruhling, H., Feussner, I., Herberg, F. W. & Maniak, M. 2013. Dictyostelium lipid droplets host novel proteins. Eukaryot Cell, 12, 1517–29.

Duckworth, B. P., Nelson, K. M. & Aldrich, C. C. 2012. Adenylating enzymes in Mycobacterium tuberculosis as drug targets. Curr Top Med Chem, 12, 766–96.

Dunn, J. D., Bosmani, C., Barisch, C., Raykov, L., Lefrancois, L. H., Cardenal-Munoz, E., Lopez-Jimenez, A. T. & Soldati, T. 2017. Eat Prey, Live: Dictyostelium discoideum As a Model for Cell-Autonomous Defenses. Front Immunol, 8, 1906.

Duvernoy, M. C., Mora, T., Ardré, M., Croquette, V., Bensimon, D., Quilliet, C., Ghigo, J. M., Balland, M., Beloin, C., Lecuyer, S. & Desprat, N. 2018. Asymmetric adhesion of rod-shaped bacteria controls microcolony morphogenesis. Nat Commun, 9, 1120.

Fishbein, S., Van Wyk, N., Warren, R. M. & Sampson, S. L. 2015. Phylogeny to function: PE/PPE protein evolution and impact on Mycobacterium tuberculosis pathogenicity. Mol Microbiol, 96, 901–16.

Franzkoch, R., Anand, A., Breitsprecher, L., Psathaki, O. E. & Barisch, C. 2024. Resolving exit strategies of mycobacteria in Dictyostelium discoideum by combining high-pressure freezing with 3D-correlative light and electron microscopy. Mol Microbiol, 121, 593–604.

Griffin, J. E., Pandey, A. K., Gilmore, S. A., Mizrahi, V., Mckinney, J. D., Bertozzi, C. R. & Sassetti, C. M. 2012. Cholesterol catabolism by Mycobacterium tuberculosis requires transcriptional and metabolic adaptations. Chem Biol, 19, 218–27.

Guallar-Garrido, S. & Soldati, T. 2024. Exploring host-pathogen interactions in the Dictyostelium discoideum-Mycobacterium marinum infection model of tuberculosis. Dis Model Mech, 17.

Hanna, N., Burdet, F., Melotti, A., Bosmani, C., Kicka, S., Hilbi, H., Cosson, P., Pagni, M. & Soldati, T. 2019. Time-resolved RNA-seq profiling of the infection of <EM>Dictyostelium discoideum</EM> by <EM>Mycobacterium marinum</EM> reveals an integrated host response to damage and stress. bioRxiv, 590810.

Hirsch, D., Stahl, A. & Lodish, H. F. 1998. A family of fatty acid transporters conserved from mycobacterium to man. Proc Natl Acad Sci U S A, 95, 8625–9.

Jones, B. S., Hu, D. D., Nicholson, K. R., Cronin, R. M., Weaver, S. D., Champion, M. M. & Champion, P. A. 2024. The loss of the PDIM/PGL virulence lipids causes differential secretion of ESX-1 substrates in Mycobacterium marinum. mSphere, 9, e0000524.

Knobloch, P., Koliwer-Brandl, H., Arnold, F. M., Hanna, N., Gonda, I., Adenau, S., Personnic, N., Barisch, C., Seeger, M. A., Soldati, T. & Hilbi, H. 2020. Mycobacterium marinum produces distinct mycobactin and carboxymycobactin siderophores to promote growth in broth and phagocytes. Cell Microbiol, 22, e13163.

Koleske, B., Shen, J., Gupta, M. & Bishai, W. R. 2025. In vivo profiling of the PE/PPE proteins of Mycobacterium tuberculosis reveals diverse contributions to virulence. Front Microbiol, 16, 1634229.

Koliwer-Brandl, H., Knobloch, P., Barisch, C., Welin, A., Hanna, N., Soldati, T. & Hilbi, H. 2019. Distinct Mycobacterium marinum phosphatases determine pathogen vacuole phosphoinositide pattern, phagosome maturation, and escape to the cytosol. Cell Microbiol, 21, e13008.

Koliwer-Brandl, H., Syson, K., Van De Weerd, R., Chandra, G., Appelmelk, B., Alber, M., Ioerger, T. R., Jacobs, W. R., JR., Geurtsen, J., Bornemann, S. & Kalscheuer, R. 2016. Metabolic Network for the Biosynthesis of Intra- and Extracellular α-Glucans Required for Virulence of Mycobacterium tuberculosis. PLoS Pathog, 12, e1005768.

Laval, T., Chaumont, L. & Demangel, C. 2021. Not too fat to fight: The emerging role of macrophage fatty acid metabolism in immunity to Mycobacterium tuberculosis. Immunol Rev, 301, 84–97.

Lee, B. S., Godejohann, M., Gürtler, F., Deloria, A. J., Liu, M., Leitgeb, R., Drexler, W., Berney, M. & Haindl, R. 2025. A functional amyloid matrix underpins the PDIM-architected corded superstructure of the <em>Mycobacterium tuberculosis</em> biofilm. bioRxiv, 2025.11.07.687260.

Li, L. O., Klett, E. L. & Coleman, R. A. 2010. Acyl-CoA synthesis, lipid metabolism and lipotoxicity. Biochim Biophys Acta, 1801, 246–51.

Lian, E., Belardinelli, J. M., De, K., Pandurangan, A. P., Angala, S. K., Palčeková, Z., Grzegorzewicz, A. E., Bryant, J. M., Blundell, T. L., Parkhill, J., Floto, R. A., Wheat, W. H. & Jackson, M. 2025. Cell envelope polysaccharide modifications alter the surface properties and interactions of Mycobacterium abscessus with innate immune cells in a morphotype-dependent manner. mBio, 16, e0032225.

Lin, H., Xing, J., Wang, H., Wang, S., Fang, R., Li, X., Li, Z. & Song, N. 2024. Roles of Lipolytic enzymes in Mycobacterium tuberculosis pathogenesis. Front Microbiol, 15, 1329715.

López-Jiménez, A. T., Cardenal-Muñoz, E., Leuba, F., Gerstenmaier, L., Barisch, C., Hagedorn, M., King, J. S. & Soldati, T. 2018. The ESCRT and autophagy machineries cooperate to repair ESX-1-dependent damage at the Mycobacterium-containing vacuole but have opposite impact on containing the infection. PLoS Pathog, 14, e1007501.

Mallick, I., Santucci, P., Poncin, I., Point, V., Kremer, L., Cavalier, J. F. & Canaan, S. 2021. Intrabacterial lipid inclusions in mycobacteria: unexpected key players in survival and pathogenesis? FEMS Microbiol Rev.

Mater, V., Eisner, S., Seidel, C. & Schneider, D. 2022. The Peripherally Membrane-attached Protein MbFACL6 of Mycobacterium tuberculosis Activates a Broad Spectrum of Substrates. J Mol Biol, 434, 167842.

Maurya, R. K., Bharti, S. & Krishnan, M. Y. 2019. Triacylglycerols: Fuelling the Hibernating Mycobacterium tuberculosis. Frontiers in cellular and infection microbiology, 8, 450–450.

Mishra, R., Hannebelle, M., Patil, V. P., Dubois, A., Garcia-Mouton, C., Kirsch, G. M., Jan, M., Sharma, K., Guex, N., Sordet-Dessimoz, J., Perez-Gil, J., Prakash, M., Knott, G. W., Dhar, N., Mckinney, J. D. & Thacker, V. V. 2023. Mechanopathology of biofilm-like Mycobacterium tuberculosis cords. Cell, 186, 5135–5150 e28.

Mohanty, D., Sankaranarayanan, R. & Gokhale, R. S. 2011. Fatty acyl-AMP ligases and polyketide synthases are unique enzymes of lipid biosynthetic machinery in Mycobacterium tuberculosis. Tuberculosis (Edinb), 91, 448–55.

Mulholland, C. V., Wiggins, T. J., Cui, J., Vilchèze, C., Rajagopalan, S., Shultis, M. W., Reyes-Fernández, E. Z., Jacobs, W. R., JR. & Berney, M. 2024. Propionate prevents loss of the PDIM virulence lipid in Mycobacterium tuberculosis. Nat Microbiol, 9, 1607–1618.

Murdoch, H. M., Dechow, S. J., Abdalla, B. J. & Abramovitch, R. B. 2026. Mycobacterium tuberculosis growth arrest on propionate at acidic pH is suppressed by mutations in phoPR and pyrazinamide treatment. mBio, 17, e0295525.

Nazarova, E. V., Montague, C. R., Huang, L., La, T., Russell, D. & Vanderven, B. C. 2019. The genetic requirements of fatty acid import by Mycobacterium tuberculosis within macrophages. Elife, 8.

Nazarova, E. V., Montague, C. R., La, T., Wilburn, K. M., Sukumar, N., Lee, W., Caldwell, S., Russell, D. G. & Vanderven, B. C. 2017. Rv3723/LucA coordinates fatty acid and cholesterol uptake in Mycobacterium tuberculosis. Elife, 6.

Orci, L., Like, A. A., Amherdt, M., Blondel, B., Kanazawa, Y., Marliss, E. B., Lambert, A. E., Wollheim, C. B. & Renold, A. E. 1973. Monolayer cell culture of neonatal rat pancreas: an ultrastructural and biochemical study of functioning endocrine cells. J Ultrastruct Res, 43, 270–97.

Padilla-Gomez, J., Mcginn, R., Liang, Y., Libby, K., Ringel, A. E., Evavold, C. & Bryson, B. D. 2025. Systems analysis reveals alternate metabolic states adopted by Mycobacterium tuberculosis across species. bioRxiv.

Pandey, A. K. & Sassetti, C. M. 2008. Mycobacterial persistence requires the utilization of host cholesterol. Proc Natl Acad Sci U S A, 105, 4376–80.

Perret, A., Bosmani, C., Leuba, F., Eblighatian, K., Guého, A., Raykov, L., Hanna, N. & Soldati, T. 2026. Membrane microdomains are crucial for Mycobacterium marinum EsxA-dependent membrane damage, escape to the cytosol, and infection. Sci Adv, 12, eady0812.

Peyron, P., Vaubourgeix, J., Poquet, Y., Levillain, F., Botanch, C., Bardou, F., Daffe, M., Emile, J. F., Marchou, B., Cardona, P. J., DE Chastellier, C. & Altare, F. 2008. Foamy macrophages from tuberculous patients’ granulomas constitute a nutrient-rich reservoir for M. tuberculosis persistence. PLoS Pathog, 4, e1000204.

Pruenster, M., Immler, R., Roth, J., Kuchler, T., Bromberger, T., Napoli, M., Nussbaumer, K., Rohwedder, I., Wackerbarth, L. M., Piantoni, C., Hennis, K., Fink, D., Kallabis, S., Schroll, T., Masgrau-Alsina, S., Budke, A., Liu, W., Vestweber, D., Wahl-Schott, C., Roth, J., Meissner, F., Moser, M., Vogl, T., Hornung, V., Broz, P. & Sperandio, M. 2023. E-selectin-mediated rapid NLRP3 inflammasome activation regulates S100A8/S100A9 release from neutrophils via transient gasdermin D pore formation. Nat Immunol, 24, 2021–2031.

Raykov, L., Mottet, M., Nitschke, J. & Soldati, T. 2023. A TRAF-like E3 ubiquitin ligase TrafE coordinates ESCRT and autophagy in endolysosomal damage response and cell-autonomous immunity to Mycobacterium marinum. Elife, 12.

Rienksma, R. A., Schaap, P. J., Martins Dos Santos, V. A. P. & SUAREZ-Diez, M. 2018. Modeling the Metabolic State of Mycobacterium tuberculosis Upon Infection. Front Cell Infect Microbiol, 8, 264.

Simwela, N. V., Jaecklein, E., Sassetti, C. M. & Russell, D. G. 2025. Impaired fatty acid import or catabolism in macrophages restricts intracellular growth of Mycobacterium tuberculosis. Elife, 13.

Tram, T. T. B., Garner, L. C., Thai, L. N. H., Nhat, L. T. H., Thu, D. D. A., Nghia, H. D. T., Van, L. H., Thwaites, G. E., Ha, V. T. N., Klenerman, P. & Thuong, N. T. T. 2025. Single-cell profiling of blood and cerebrospinal fluid in tuberculous meningitis. J Immunol, 214, 2894–2905.

Trivedi, O. A., Arora, P., Sridharan, V., Tickoo, R., Mohanty, D. & Gokhale, R. S. 2004. Enzymic activation and transfer of fatty acids as acyl-adenylates in mycobacteria. Nature, 428, 441–5.

Trofimov, V., Kicka, S., Mucaria, S., Hanna, N., Ramon-Olayo, F., Peral, L. V., Lelièvre, J., Ballell, L., Scapozza, L., Besra, G. S., Cox, J. A. G. & Soldati, T. 2018. Antimycobacterial drug discovery using Mycobacteria-infected amoebae identifies anti-infectives and new molecular targets. Sci Rep, 8, 3939.

Vanderven, B. C., Fahey, R. J., Lee, W., Liu, Y., Abramovitch, R. B., Memmott, C., Crowe, A. M., Eltis, L. D., Perola, E., Deininger, D. D., Wang, T., Locher, C. P. & Russell, D. G. 2015. Novel inhibitors of cholesterol degradation in Mycobacterium tuberculosis reveal how the bacterium’s metabolism is constrained by the intracellular environment. PLoS Pathog, 11, e1004679.

Wang, Q., Boshoff, H. I. M., Harrison, J. R., Ray, P. C., Green, S. R., Wyatt, P. G. & Barry, C. E., 3RD 2020. PE/PPE proteins mediate nutrient transport across the outer membrane of Mycobacterium tuberculosis. Science, 367, 1147–1151.

Weaver, S. D., Hu, D. D., Jones, B. S., Champion, P. A. & Champion, M. M. 2025. PDIM/PGL-deficient Mycobacterium marinum shows disrupted protein secretion measured with bottom-up label-free quantitative analysis in data-dependent acquisition mode on a timsTOF Pro 2 mass spectrometer. *Microbiol Resour Announc*, e0116125.

Weimar, J. D., Dirusso, C. C., Delio, R. & Black, P. N. 2002. Functional role of fatty acyl-coenzyme A synthetase in the transmembrane movement and activation of exogenous long-chain fatty acids. Amino acid residues within the ATP/AMP signature motif of Escherichia coli FadD are required for enzyme activity and fatty acid transport. J Biol Chem, 277, 29369–76.

Wilburn, K. M., Fieweger, R. A. & Vanderven, B. C. 2018. Cholesterol and fatty acids grease the wheels of Mycobacterium tuberculosis pathogenesis. Pathog Dis, 76.

