## Supplementary figures and images for "Intracellular compartmentalization shapes lipid access and metabolic fitness of mycobacteria"

### Supplementary figures

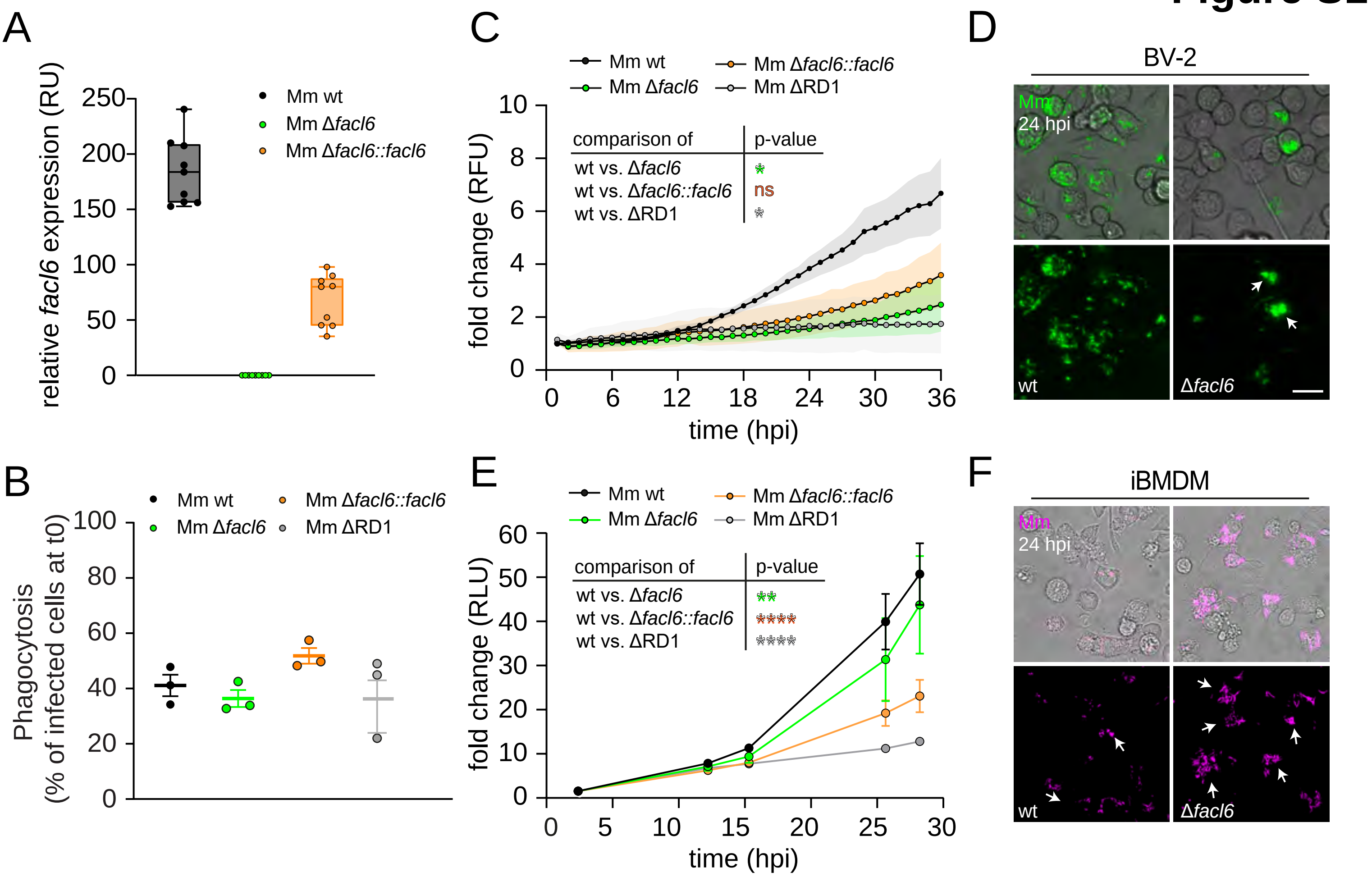

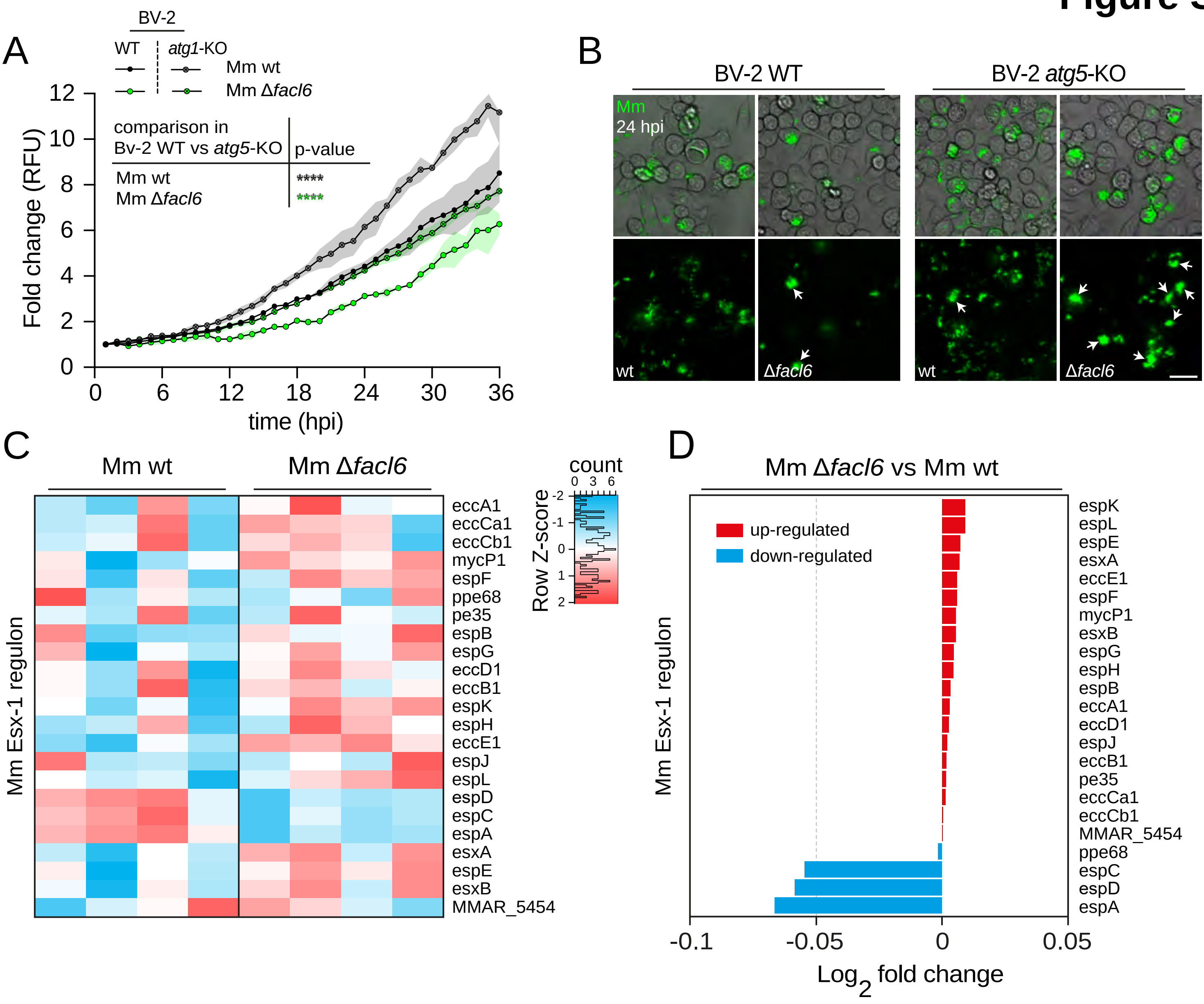

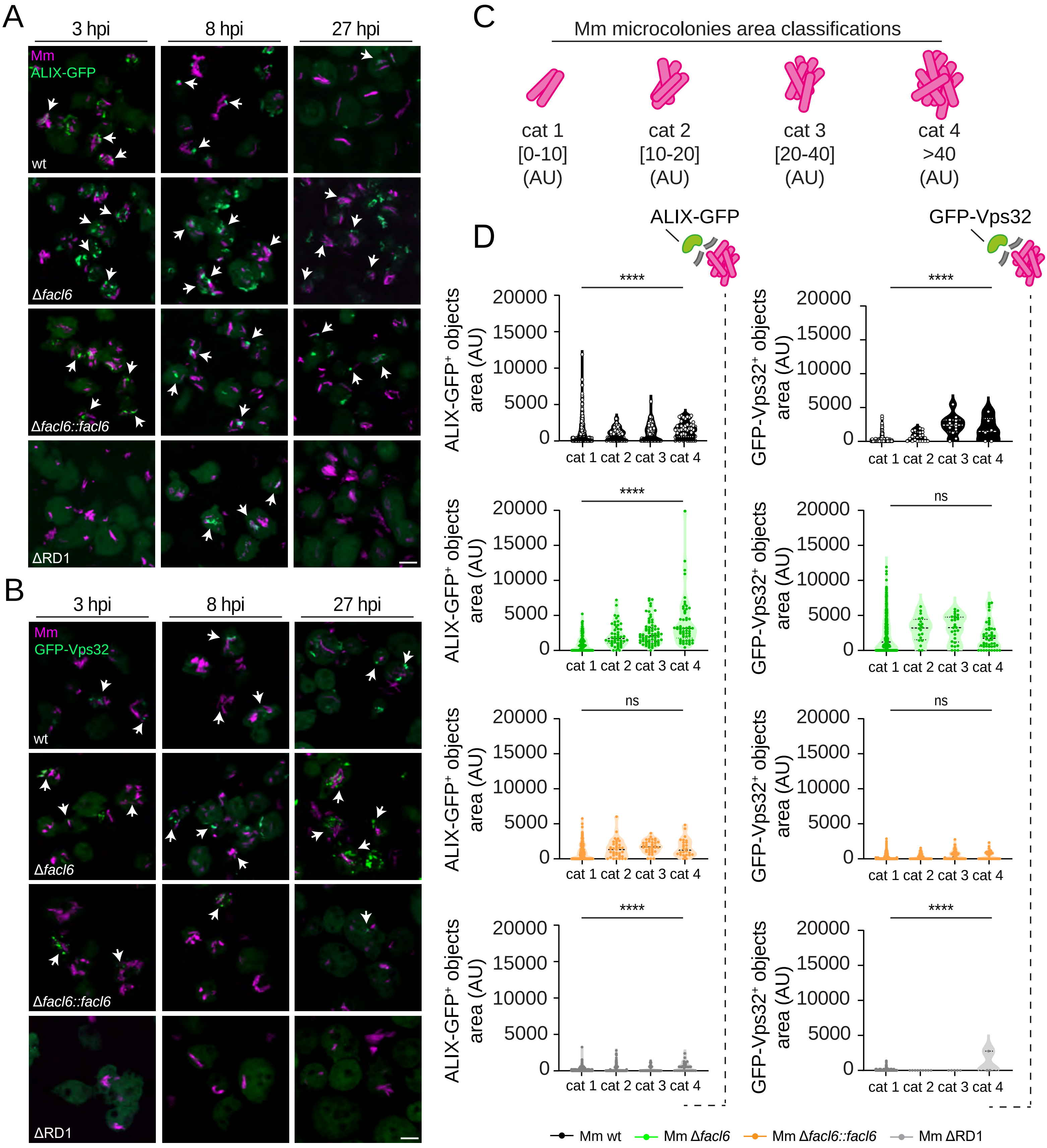

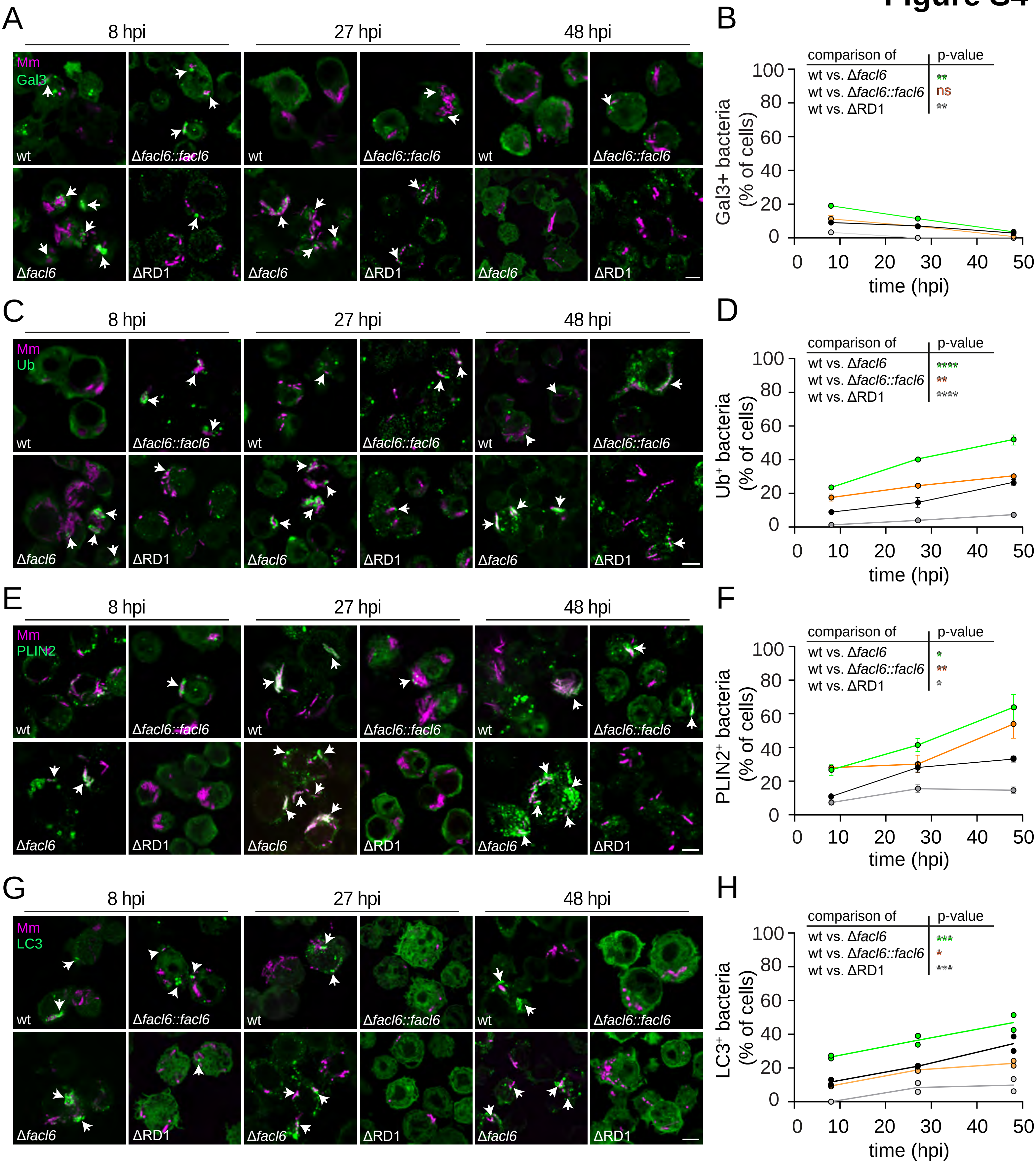

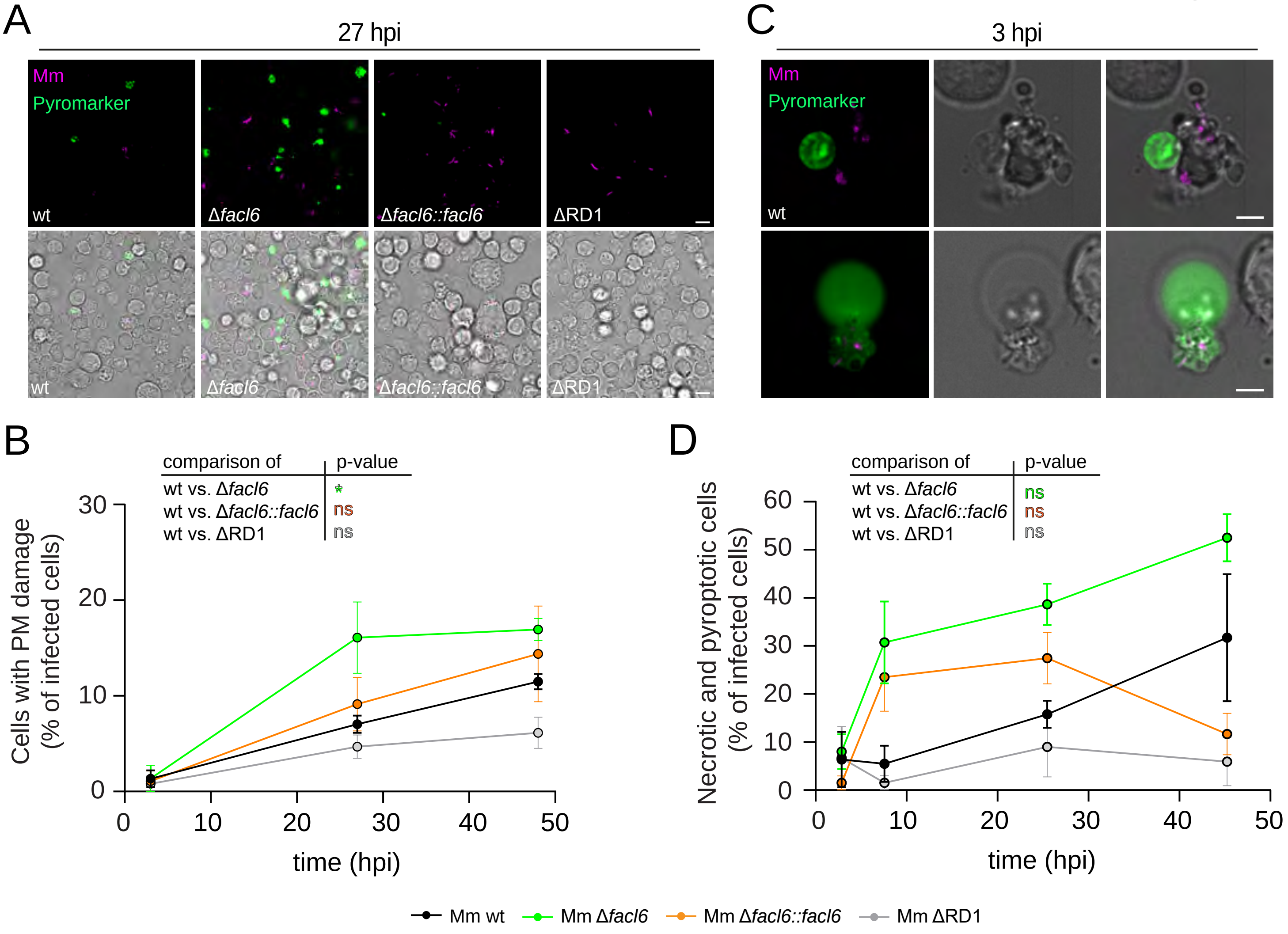

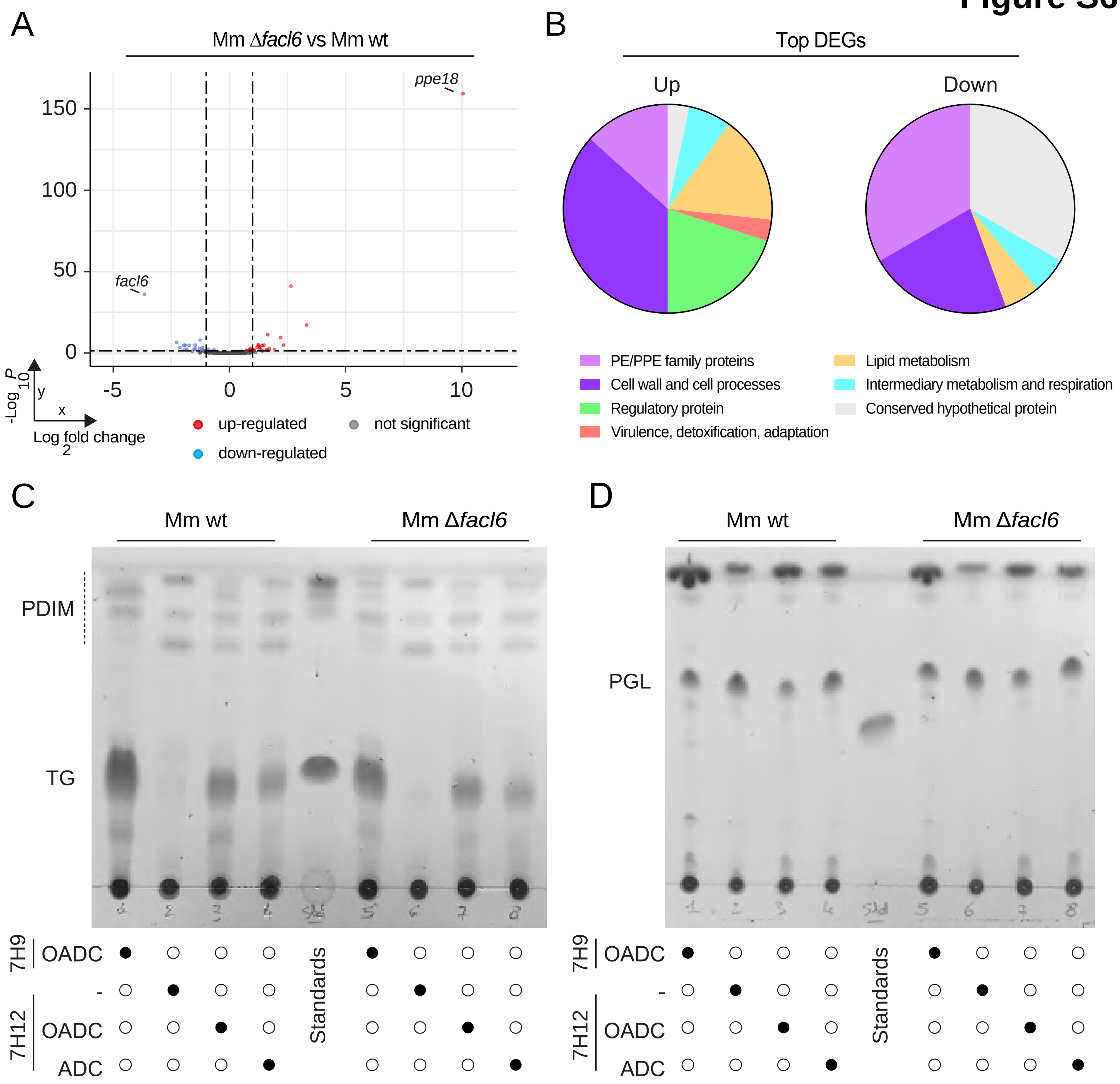

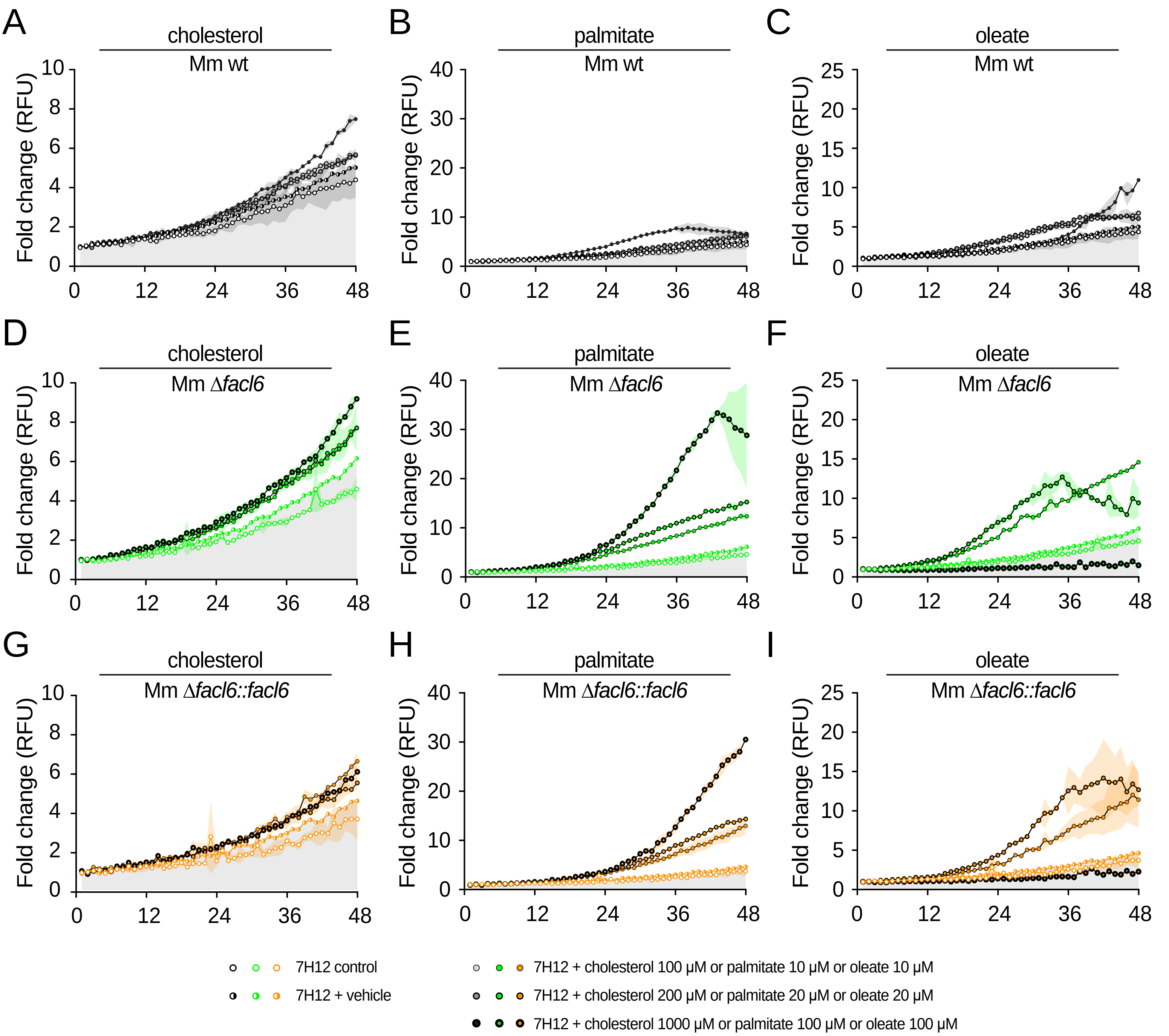
